# Precise detection of S phase onset reveals decoupled G1/S transition events

**DOI:** 10.1101/300442

**Authors:** Gavin D. Grant, Katarzyna M. Kedziora, Juanita C. Limas, Jeremy E. Purvis, Jeanette Gowen Cook

## Abstract

The eukaryotic cell division cycle is the process by which cells duplicate their genomes and proliferate. Transitions between sequential cell cycle phases are tightly orchestrated to ensure precise and efficient cell cycle progression. Interrogating molecular events at these transitions is important for understanding normal and pathological cell proliferation and mechanisms that ensure genome stability. A popular fluorescent reporter system known as “FUCCI” has been widely adopted for identifying cell cycle phases. Using time-lapse fluorescence microscopy, we quantitatively analyzed the dynamics of the FUCCI reporters relative to the transitions into and out of S phase. Although the original reporters reflect the E3 ubiquitin ligase activities for which they were designed, SCF^Skp2^ and APC^Cdh1^, their dynamics are significantly and variably offset from actual S phase boundaries. To precisely mark these transitions, we generated and thoroughly validated a new reporter containing a <u>P</u>CNA-<u>i</u>nteracting <u>p</u>rotein degron whose oscillations are directly coupled to the process of DNA replication itself. We combined this reporter with the geminin-based APC^Cdh1^ reporter to create “PIP-FUCCI.” PIP degron reporter dynamics closely correlate with S phase transitions irrespective of reporter expression levels. Using PIP-FUCCI, we made the unexpected observation that the apparent timing of APC^Cdh1^ inactivation frequently varies relative to the onset of S phase. We demonstrate that APC^Cdh1^ inactivation is not a strict pre-requisite for S phase entry, though delayed APC^Cdh1^ inactivation correlates with longer S phase. Our results illustrate the benefits of precise delineation of cell cycle phase boundaries for uncovering the sequences of molecular events at critical cell cycle transitions.

## Introduction

The eukaryotic cell division cycle is tightly regulated to ensure timely cell proliferation while maintaining genome stability. For this reason, monitoring progression through the sequential cell cycle phases is an important component of many biological studies. A variety of quantitative tools are in wide use to assess cell cycle phase position (analytical flow cytometry, antibody markers of cell cycle proteins, DNA stains, etc.), but most require harvesting cells at a single time point and thus provide little or no information about cell cycle dynamics. Until recently, most cell cycle dynamics information has come from arresting cell populations at a single cell cycle phase and collecting time points after release from the cell cycle block. The synchronization methods themselves introduce strong perturbations and cellular stresses, which are not well tolerated by all cell lines, and these perturbations can confound interpretation even in those cells that are synchronized. Moreover, intercellular heterogeneity after release from a block leads to rapid loss of synchrony in nearly all mammalian cell lines, further blurring the dynamics of cell cycle progression.

To address these limitations, automated time-lapse live imaging of actively proliferating cells has become an increasingly popular method of studying cell cycle dynamics. This approach is particularly powerful because the time resolution can be high (minutes or even seconds) relative to cell cycle length (hours or days), and the pitfalls of artificial synchronization are bypassed. Cell timelines can be readily synchronized *in silico* to derive a statistical average of the population (1). Importantly, analyzing single cells reveals unique behaviors in subpopulations that would be missed by ensemble assays such as immunoblots or proteomics (2, 3). An important advance for metazoan cell cycle investigations was the introduction of a pair of genetically encoded fluorescent reporters, known as the Fluorescent, Ubiquitination-based Cell Cycle Indicator, or FUCCI (4). FUCCI reporters are based on two fragments of human DNA replication origin licensing proteins, Cdt1 and geminin, fused to different fluorescent proteins. One fragment is from human Cdt1 (Cdt1_30-120_), which is stable during G1 and early S phase and ubiquitylated by SCF^Skp2^ from ~mid-S phase through mitosis (5). The second fragment contains a portion of the human origin licensing inhibitor, Geminin (Gem_1-110_), which is ubiquitylated from anaphase through G1 by APC^Cdh1^ (6). These two reporters were designed to have roughly reciprocal accumulation and degradation patterns over the full duration of the cell cycle, and they have been widely adopted and adapted to different applications. FUCCI derivatives include versions with different fluorophore tags, encode the two reporters from a single expression vector, or have been optimized for expression in different species (7–11). Despite their popularity, a detailed quantitative analysis of the original FUCCI reporters has not yet been reported.

The original FUCCI reporters are excellent tools for approximating cell cycle phase transitions and measuring the timing of cell proliferation. They do not precisely mark either the G1/S or the S/G2 transition, however. For example, we demonstrate here that the FUCCI reporters routinely mis-identify S phase cells as G1 cells. This misidentification is meaningful because the onset of S phase marks a critical and generally irreversible cellular commitment to proliferation. Moreover, the original study reported the unexpected difficulty of using different colored fluorescent proteins for the Cdt1 reporter: only one color successfully oscillated as expected (4, 11). Sakaue-Sawano and colleagues also recently described a new derivative FUCCI reporter pair that responds more robustly to S-phase-specific ubiquitylation and degradation, underscoring the need for fully validated tools to precisely mark the G1/S transition (12). We hypothesized that close examination of dynamics at the G1/S transition will reveal new aspects of this critical cell cycle event that have important consequences for understanding cell proliferation and genome maintenance.

In this study, we sought to fully understand the temporal dynamics of the FUCCI reporters, to identify their advantages and drawbacks, and to provide a mechanistic explanation for the counter-intuitive behavior of the original FUCCI Cdt1-based reporter in particular. Toward this end, we have designed and fully validated an improved and robust set of reporters that very precisely mark the G1/S transition in multiple experimental settings. The reporter is based on a PCNA-interacting protein motif (PIP) degron in combination with the geminin-based APC^Cdh1^ reporter, or “PIP-FUCCI”. Using PIP-FUCCI in combination with the DNA replication reporter, PCNA (proliferating cell nuclear antigen), we uncovered surprisingly variable differences between S phase onset and the apparent timing of APC^Cdh1^ inactivation. We provide guidance to investigators in selecting appropriate fluorescent cell cycle reporters and new analysis strategies for delineating cell cycle transitions. These findings illustrate the utility of precise cell cycle transition reporters and contribute to ongoing investigations of cell cycle dynamics and the sequence of events at the critical G1/S transition.

## Results

### FUCCI dynamics are offset from the G1/S phase transition

Our first goal was to comprehensively define the dynamics of the original FUCCI reporters relative to the boundaries of S phase. To define those boundaries, we stably expressed the full-length PCNA protein fused to mTurquoise2 (mTurq2-PCNA) in a population of U2OS cells (human osteosarcoma) as it is an established localization-based reporter of S phase progression (13–15). We expressed the PCNA fusion at a low level (Fig. S1A) and observed no deleterious effects both in standard culture and under imaging conditions (unpublished observations). Images from each hour of a cell cycle of one cell expressing mTurq2-PCNA are shown in Fig. 1A (top row). During S phase, PCNA is loaded at DNA replication forks forming foci in stereotypical punctate patterns that change from early S to late S phase (13, 16, 17). PCNA localization is precisely correlated with DNA replication and thus is a *bona fide* marker of S phase (13-16, 18, 19). At DNA replication onset, PCNA is loaded onto DNA in small foci that are evenly distributed in the nucleus. By late S phase, PCNA localizes to a few large foci. During both G1 and G2 phases, PCNA is nuclear but diffuse because it is not DNA-loaded. At the G2/M transition after nuclear envelope breakdown, PCNA is diffuse throughout the cells. The transition from diffuse to punctate and from punctate back to diffuse marks the G1/S and S/G2 transitions respectively, and these changes are often visible between two consecutive frames of the video (i.e. within 10 minutes).

**Figure 1.**
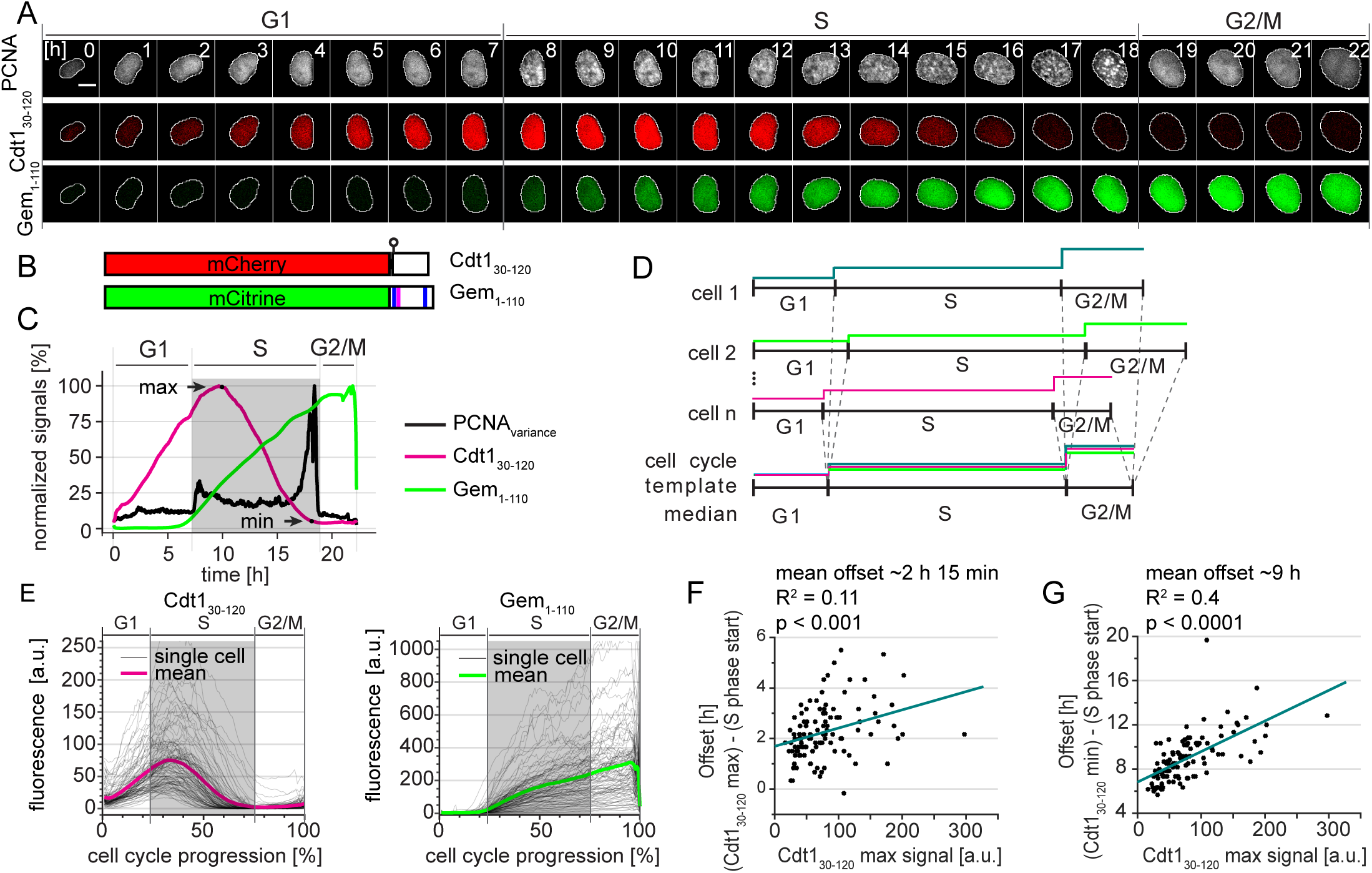
FUCCI dynamics are offset from S phase. A. Selected images from wide field time-lapse imaging of a U2OS cell expressing PCNA-mTurq2 and FUCCI fragments Cdt1_30-120_-mCherry and Gem*1-110*-Citrine. One frame from each hour beginning after cytokinesis is shown with the nucleus outlined; numbers in the upper strip indicate hours since mitosis. Scale bar 10 μm.
B. Schematic of the two fluorescent fragments used in FUCCI reporters from Sakaue-Sawanao 2008 and closely related derivatives (4). The human Cdt1 S31 phosphorylation site is indicated by a circle. Human geminin motifs for targeting by APC^Cdh1^, D box (magenta line), and KEN motifs (blue lines) are marked.
C. Quantification of fluorescence signals from the single cell presented in 1A. Mean fluorescence intensity of FUCCI fragments within nuclear regions were normalized to their maximum values. The PCNA signal variance produced by PCNA loading into nuclear foci was analyzed as described in Materials and Methods; the gray rectangle marks S phase. Images were collected every 10 minutes.
D. Illustration of the time warping method to align many traces from individual cells for inspection of cell cycle phase transitions (for details see Materials and Methods).
E. Single cell traces (gray lines) and mean (magenta and green) of FUCCI reporter dynamics in individual U2OS cells of a clonal population. All traces were warped as in 1D, the total time between cell divisions is set to one hundred percent, and gray rectangles show the median S phase period of the population based on PCNA foci. Cells were imaged every 10 min (n=107).
F. The correlation of the Cdt1_30-120_ maximum signal (see 1C) and the time delay between S phase onset determined by PCNA foci and the time of Cdt1_30-120_ peak in individual cells in 1E. The teal line represents the best fit linear model.
G. The correlation of the Cdt1_30-120_ maximum signal (see 1C) and the time delay between S phase onset determined by PCNA foci and the disappearance of Cdt1_30-120_ signal (as indicated by Cdt1 min in 1C) in individual cells in 1E. The teal line represents the best fit linear model.

We analyzed FUCCI reporters - Cdt1_30-120_ expressed as a fusion to mCherry and Gem_1-110_ expressed as a fusion to Citrine (Fig. 1B) in the same cells as mTurq2-PCNA (Fig. 1A). We selected clonal populations expressing all three reporters. We subjected these cells to time-lapse fluorescence imaging over the course of 72 hours of unperturbed proliferation and collected images every 10 minutes. We detected no significant changes in cell cycle duration among cells analyzed early or late in the imaging period (Fig. S1B). We employed semi-automated segmentation and tracking tools to efficiently generate many fluorescence intensity traces from each experiment for rigorous statistical analysis.

To mark S phase boundaries, we developed a sensitive approach based on calculating the spatially-localized variance of PCNA intensity across the nucleus (details of analysis in Materials and Methods section). This variance is quantified over time by the black line in Fig. 1C with time zero set to the first frame after cytokinesis. The rapid increase in PCNA variance indicates the onset of DNA replication whereas the steep drop in PCNA variance indicates the end of S phase. Thus, we can precisely define the temporal position of any given cell within interphase. For display purposes we use gray shading to indicate S phase duration calculated by PCNA variance here and in subsequent figures (Fig. 1C).

We next sought to quantify the timing of the FUCCI system relative to PCNA-based cell cycle phase detection. We observed that peak expression of mCherry-Cdt1_30-120_ occurred well *after* Citrine-Gem_1-110_ began to accumulate and PCNA had formed puncta. Traces of these three reporters from a single cell are shown in Figure 1C. The increase in Citrine-Gem_1-110_ accumulation closely coincided with the onset of S phase in this U2OS cell, but the degradation of mCherry-Cdt1_30-120_ was delayed. This overlap of the Cdt1-based reporter with early S phase is consistent with the detection of BrdU-incorporation in Cdt1_30-120_-expressing cells shown in Sakaue-Sawano (4). Moreover, mCherry-Cdt1_30-120_ was still present even into late S phase (Fig. 1A and C).

Because the durations of cell cycle phases vary between single cells (3, 20–25), to visualize FUCCI reporters dynamics in a population of cells we developed a computational strategy to adjust for cell-to-cell variation in phase duration. Using PCNA variance-based phase boundaries, we independently interpolated individual traces *within* each G1, S phase, and G2/M period. We adjusted these three portions for each trace to the length of the median fraction of total cell cycle duration in the cell population (G1: 22%, S: 53%, and G2/M: 25% in this population of U2OS cells, Fig. 1D). This phase-specific “time warping” provides enhanced time resolution compared to aligning only to a single event (e.g. cytokinesis) and accentuates rapid events that occur near phase transitions. We did not observe any artefacts near the phase transitions from applying this method to an unregulated nuclear reporter, NLS-Venus (Fig. S1C). Using this approach, we observed that the onset of Citrine-Gem_1-110_ accumulation began consistently close to the beginning of S phase whereas the substantial delay in the peak of mCherry-Cdt1_30-120_ relative to S phase onset was apparent in the majority of U2OS cells (Fig. 1E). Interestingly, the offset between the start of S phase and peak mCherry-Cdt1_30-120_ intensity was correlated with the expression level of the Cdt1_30-120_ reporter (Fig. 1F). This offset reveals an important caveat for using peak Cdt1_30-120_ expression to mark the timing of the G1/S transition: marking the G1/S transition will depend on how highly the reporter is expressed in a given cell. Similarly, the timing of complete mCherry-Cdt1_30-120_ degradation also correlated with the maximum intensity expression (Fig 1G). These results were reproducible in an independent U2OS clone with particularly high reporter expression (Supplementary Fig. S2).

### FUCCI dynamics rely on a phosphomimetic linker

An important strength of the FUCCI reporter system is the use of inert protein fragments of both Cdt1 and geminin. Interestingly, the Cdt1 fragment, which contains amino acids 30-120, interrupts the CDK-primary phosphorylation site for SCF^Skp2^ recognition, which is threonine-proline at positions 29-30 in the native protein. A second CDK site is serine-proline at 31-32, but prior work indicated that this site is a weak target for SCF^Skp^ binding (26, 27). We therefore considered the possibility that the offset of Cdt_130-120_ degradation occurs because the major SCK^Skp2^ binding site at T29 is missing.

To test if a reporter containing the complete CDK-dependent phosphodegron marks the onset of S phase more precisely, we expanded the Cdt1 fragment to include amino acids 16 to 120 (Fig. 2A). We stably expressed mCherry-Cdt1_16-120_ in U2OS cells and monitored its dynamics relative to the S phase boundaries defined by mTurq2-PCNA as in Fig. 1E. Despite the addition of the primary phosphorylation site at T29, the mCherry-Cdt1_16-120_ fragment accumulated and degraded with similar kinetics as FUCCI mCherry-Cdt1_30-120_ (Fig. 2B). Upon closer inspection of the sequences of the original FUCCI construct, we noted the presence of a glutamate in the linker connecting Cdt1_30-120_ to the fluorescent protein in both the original construct and a derivative for expression in stem cells (ES-FUCCI) (4, 8). Since glutamate can sometimes mimic constitutive phosphorylation, we hypothesized that the glutamate residue at this position analogous to T29 in native Cdt1 mimics Cdt1 phosphorylation for SCF^Skp2^ targeting. To test this idea directly, we constructed two variants of the linker with either alanine (E29A) or threonine (E29T) at the final position of the linker which corresponds to position 29 (e.g. threonine) in full length Cdt1 (Fig 2A). Cdt1 oscillations were virtually abolished in the E29A variant (Fig. 2C), suggesting that S31 is either not phosphorylated or, as in native Cdt1, is not an effective Skp2 binding site. On the other hand, the E29T variant phenocopied the original Cdt1_30-120_ reporter containing glutamate in its linker (Fig. 2D). We conclude that the original Cdt1_30-120_ FUCCI fragment oscillates because of a phosphomimetic amino acid in the linker region that compensates for the absence of the original phosphothreonine in native Cdt1. Moreover, since E29T, the original FUCCI fragment and linker (E29), and the Cdt1_16-120_ constructs all have similar cell cycle kinetics, we postulate that they are approximately equivalent SCF^Skp2^ activity sensors in these cells. Moreover, the E29 version is a strict SCF^Skp2^ activity biosensor that is independent of substrate phosphorylation.

**Figure 2.**
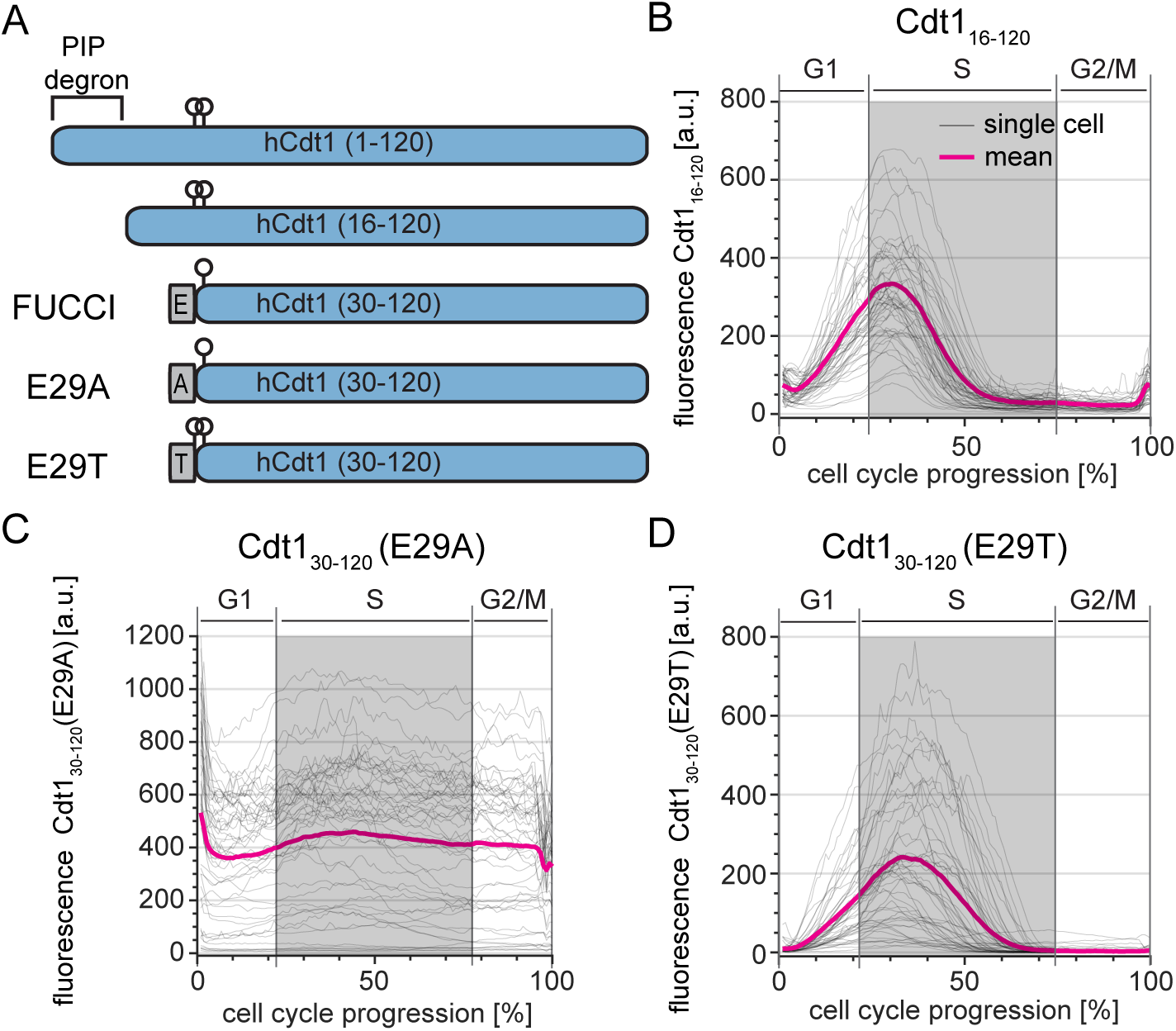
FUCCI Cdt1_30-120_ is a phosphomimetic SCF^Skp2^ sensor. A. Schematics of the first 120 amino acids of Cdt1 and the four different reporter fragments analyzed in this study. Relevant phosphorylation sites are indicated by circles; linker regions derived from expression vectors are gray rectangles.
B. Dynamics of the Cdt1_16-120_-mCherry reporter throughout the cell cycle analyzed as in Fig. 1E. Gray lines are 50 randomly selected cells and the magenta line is the mean signal for the population (n=52).
C. Dynamics of the Cdt1_30-120_ (E29A)-mCherry reporter (50 random traces (gray) and mean of the population (magenta), n=107).
D. Dynamics of the Cdt1_30-120_ (E29T)-mCherry reporter (50 random traces (gray) and mean of the population (magenta), n=73).

### A robust degradation-based S phase sensor

Although the PCNA reporter precisely marks the boundaries of S phase, its use requires high-resolution microscopy and image processing that is more complex than simple fluorescence intensity measurements. The simplicity of the degradation-based FUCCI system is an attractive practical feature that we sought to optimize. For this purpose, we designed a reporter that is subject to replication-coupled destruction by CRL4^Cdt2^. Many substrates of the CRL4^Cdt2^ E3 ubiquitin ligase are only recognized for ubiquitylation when bound to DNA-loaded PCNA. This requirement for interaction with loaded PCNA tightly couples substrate degradation to DNA replication because PCNA is loaded as an integral component of replication forks in S phase. We generated a minimal expression construct consisting of the PCNA-interacting protein (PIP) degron from human Cdt1 (Cdt1_1-17_), fused to a nuclear localization sequence, an HA tag, and the mVenus fluorescent protein; we named this fusion “PIP-mVenus”. The Cdt1 PIP degron has been well-characterized both in native Cdt1 and as a fusion to heterologous proteins. Fusing the Cdt1 PIP degron to heterologous proteins confers particularly rapid replication-coupled degradation compared to the PIP degron from another CRL4^Cdt2^ substrate, the CDK inhibitor p21 (28). We expressed this replication sensor with a P2A “self-cleaving” peptide linker to the same mCherry-Gem_1-110_ from the FUCCI system (29). We refer to this construct (PIP-mVenus-P2A-mCherry-Gem_1-110_) as “PIP-FUCCI” (PIP-Fluorescent Ubiquitylation Cell Cycle Indicator) (Fig. 3A).

**Figure 3.**
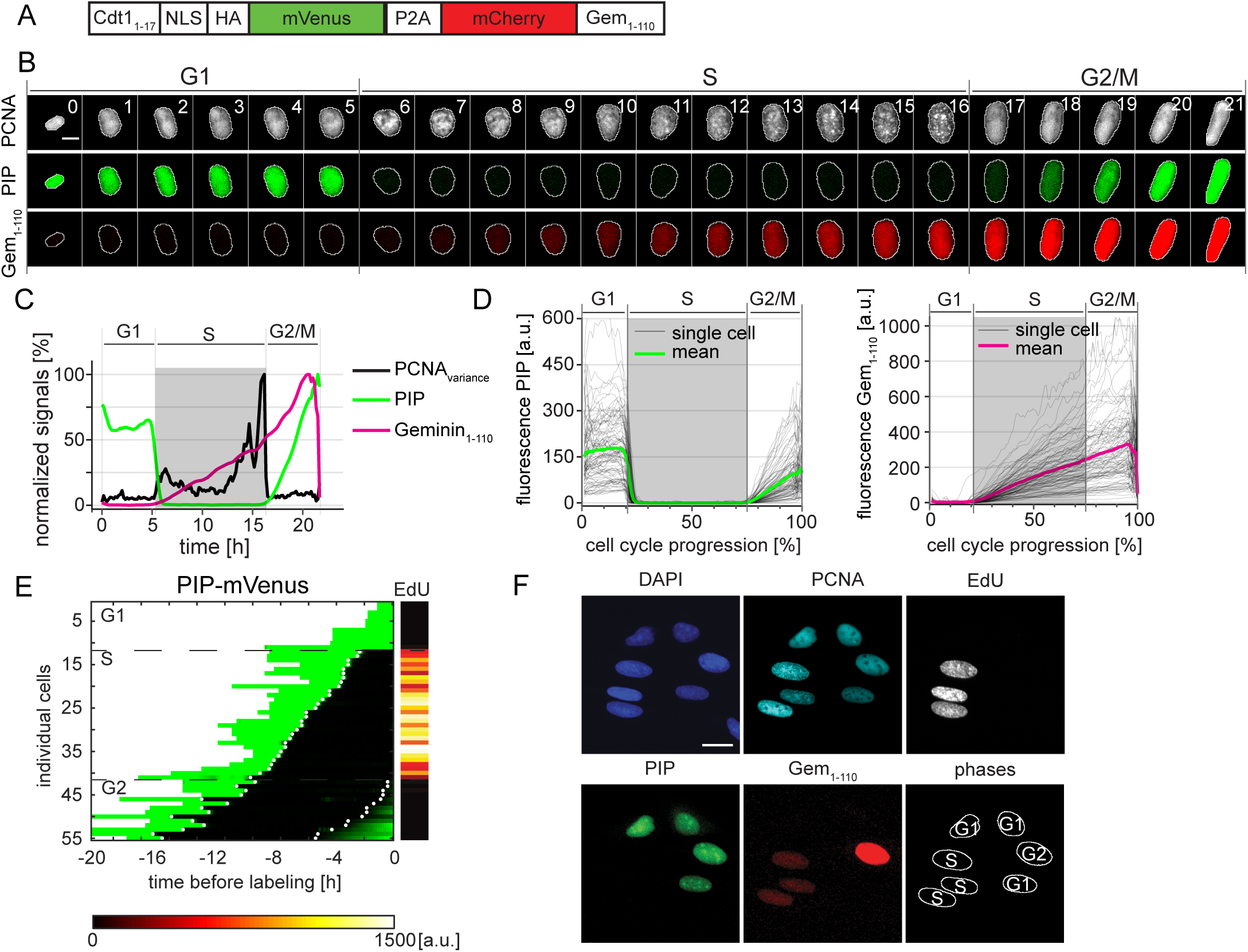
PIP-FUCCI is a precise cell cycle phase indicator. A. Schematic of the PIP-FUCCI dual reporter expression construct. Cdt1_1-17_ = human Cdt1 amino acids 1-17 including the PIP degron, NLS= SV40 nuclear localization signal, HA=epitope tag, P2A= self-cleaving peptide, Gem_1-110_= human Geminin amino acids 1-110 including both the D box and KEN motif.
B. Selected images from wide field time-lapse imaging of a U2OS cell expressing PCNA-mTurq2 and the PIP-FUCCI reporters PIP-mVenus and mCherry-Gem_1-110_. One frame from each hour beginning after cytokinesis is shown with the nucleus outlined; numbers in the upper strip indicate hours since mitosis. Scale bar 10 μm.
C. Quantification of PCNA variance (black line) and PIP-FUCCI reporter fluorescence signals (green and red lines) from the cell in 3B.
D. Dynamics of the PIP-mVenus reporter (left) and the mCherry-Gem_1-110_ (right) throughout the cell cycle. Gray lines are 100 randomly selected cells and heavy colored lines are the mean signal in the population (n=125). S phase boundaries were determined by PCNA localization and are indicated with a gray rectangle; warping as in figure 1D.
E. Heat map of the signal intensity values of PIP-mVenus (green) in 55 asynchronously growing cells before EdU labeling. EdU intensity value (right) of the 55 cells. White dots indicate the beginning and end of S phase annotated manually based on the PCNA signal.
F. Micrographs of selected asynchronously growing U2OS cells before EdU labeling (PCNA, PIP, and Gem_1-110_) and after labeling (DAPI and EdU). The cell cycle phase of each cell is indicated in the bottom right panel. Scale bar, 25 μm.

We stably expressed this two-reporter single construct in U2OS cells with the PCNA reporter and collected time-lapse images as before. Similar to an analogous “FUCCI(CA)” construct (12), PIP-mVenus fluorescence was rapidly lost at the onset of DNA replication concurrent with PCNA foci formation (Fig. 3B-D and Supplementary Movie 1). At the end of S phase, PIP-mVenus accumulated throughout G2 in an almost linear fashion. The changes in PIP-FUCCI protein dynamics at the G1 to S phase transition and the S phase to G2 transition closely corresponded with the changes in PCNA variance associated with DNA replication. To test if the new PIP-mVenus fusion is strictly absent throughout all of S phase, we imaged cells every 10 minutes for 22 hours followed by a pulse of EdU for 30 minutes. After processing for EdU detection, we matched cells to their time lapse images. Unlike the reporters based on Skp2 targeting, 100% of PIP-mVenus positive cells were EdU-negative with diffuse PCNA localization, whereas all EdU-positive cells were PIP-mVenus-negative with focal PCNA localization (Fig. 3E).

The remarkably tight correlation between PIP-mVenus degradation and the entirety of S phase combined with the restriction of mCherry-Gem_1-110_ expression to S phase and G2 provides a means to unambiguously identify cell cycle phases in fixed cells. To illustrate this use, we analyzed cells by PIP-mVenus and mCherry-Gem_1-110_ expression and secondarily by PCNA localization and EdU incorporation (Fig. 3F). We readily identified G1 cells as mVenus single-positive, S phase cells as mCherry single positive, G2 cells as nuclear mVenus and mCherry double-positives; and M phase cells had both fluorescent proteins diffusely expressed since their nuclear envelopes had been disassembled. Thus, just two reporters (PIP-mVenus and mCherry-Gem_1-110_) are sufficient to accurately identify cell cycle phases in fixed cells.

We also performed a detailed analysis of the reliability for calling cell cycle phases in time lapse experiments using only the intensity levels of PIP-mVenus and compared this approach to the more demanding analysis of PCNA variance. We defined the start of S phase as the point when the intensity value of PIP-mVenus was half of its maximal value in G1; we call this point “PIP_mid_” (Fig. 4A); we note that this point is a few minutes after the true S phase onset given the requirement for PCNA loading before PIP-mVenus degradation. When using PIP-mVenus as an indicator of the S to G2 transition we use the onset of signal accumulation (termed “PIP_rise_”). PIP-mid is tightly correlated with the onset of PCNA foci formation (quantified by the increase in PCNA variance) with an offset of approximately 5 minutes. Importantly, unlike the S phase offset in Cdt1_30-120_ degradation, different PIP-mVenus reporter expression levels had no effect on degradation timing relative to S phase in the studied population; even very high reporter expression accurately marks S phase onset (Fig. 4B p=0.26). To further define the accuracy of PIP-FUCCI as a cell cycle reporter, we compared cell cycle phase durations defined by PCNA variance or PIP-FUCCI intensity levels in both U2OS cells and in non-transformed RPE-hTert cells (Supplementary Movie 2). We found PIP-FUCCI to be just as accurate as PCNA variance for defining cell cycle phases in both cell lines (Fig. 4C).

**Figure 4.**
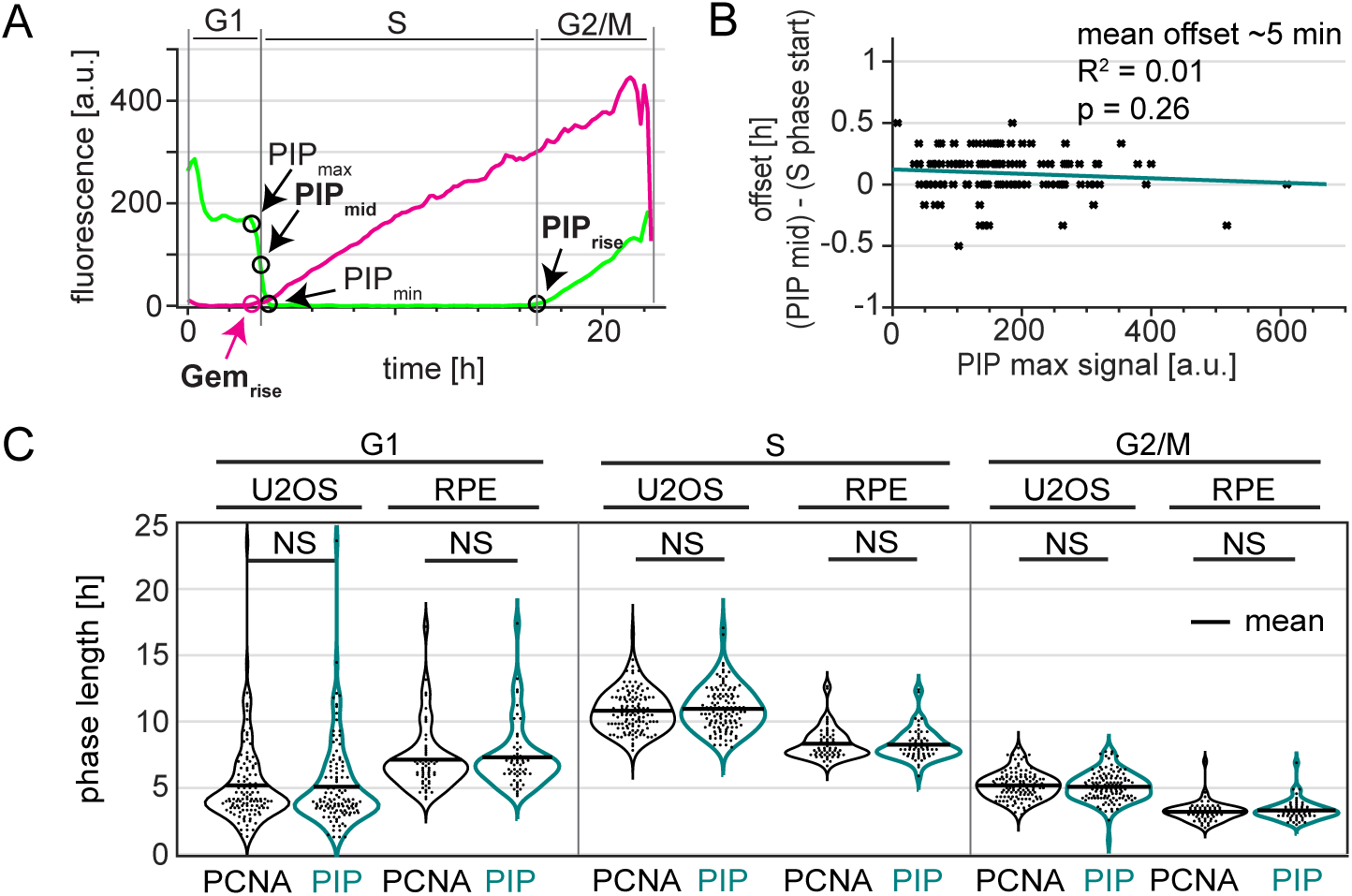
The PIP degron fusion protein precisely marks the G1/S and S/G2 transitions. A. One annotated example of traces from a single U2OS cell expressing PIP-mVenus (green) and mCherry-Gem_1-110_ (red). “PIP_mid_” is the time at which 50% of the maximum PIP-mVenus signal is lost ((“PIP_max_” – “PIP_min_”)/2) is lost and is used to mark the beginning of S phase. Details of trace annotations are described in Materials and Methods.
B. Correlation of PIP-mVenus PIP_max_ level with the offset between the beginning of S phase (determined by PCNA foci) and PIP_mid_ in individual U2OS cells. Teal line represents the best fit linear model (n=121).
C. Distributions of cell cycle phase lengths measured by PCNA foci and PIP-mVenus dynamics in U2OS (n=121) and RPE-hTert (n=57) cells. No differences between the pairs of distributions were detected according to Kolmogorov-Smirnov test (p>0.05). Black line indicates mean.

### The effects of DNA damage on PIP-FUCCI dynamics

PCNA is also loaded during DNA damage repair. This PCNA loading is a prerequisite for substrate interactions with the CRL4^Cdt2^ E3 ubiquitin ligase, which targets PIP-degron containing substrates (28, 30–39). CRL4^Cdt2^-mediated substrate degradation has different kinetics based on the cell cycle timing of the DNA insult: degradation is efficiently stimulated by DNA damage in G1, is inefficient in G2 cells due to CDK1-mediated CRL4^Cdt2^ inactivation, and DNA damage induces little additional substrate degradation in S phase since PCNA is already loaded during DNA replication. (40, 41). To define the dynamics of PIP-FUCCI after DNA damage, we treated PIP-FUCCI U2OS cells with a high dose (200 ng/mL) of the radiomimetic DNA damaging agent neocarzinostatin (NCS) (Fig. 5A and 5B). We imaged cells for approximately 24 hours before NCS treatment to establish a precise cell cycle position at the time of damage. As expected, NCS-treated U2OS cells in G1 rapidly lost PIP-mVenus signal followed by gradual PIP-mVenus re-accumulation (Fig 5A top left panel). In contrast, G2 cells showed only moderate reduction in PIP-mVenus signal intensity upon NCS treatment (Fig. 5A top right panel and 5B). PIP-mVenus was undetectable in S phase (Fig.5Ci and unpublished observations) and remained so after NCS treatment. Thus, PIP-mVenus shows a rapid decrease in signal in response to large doses of DNA damage only during G1.

**Figure 5.**
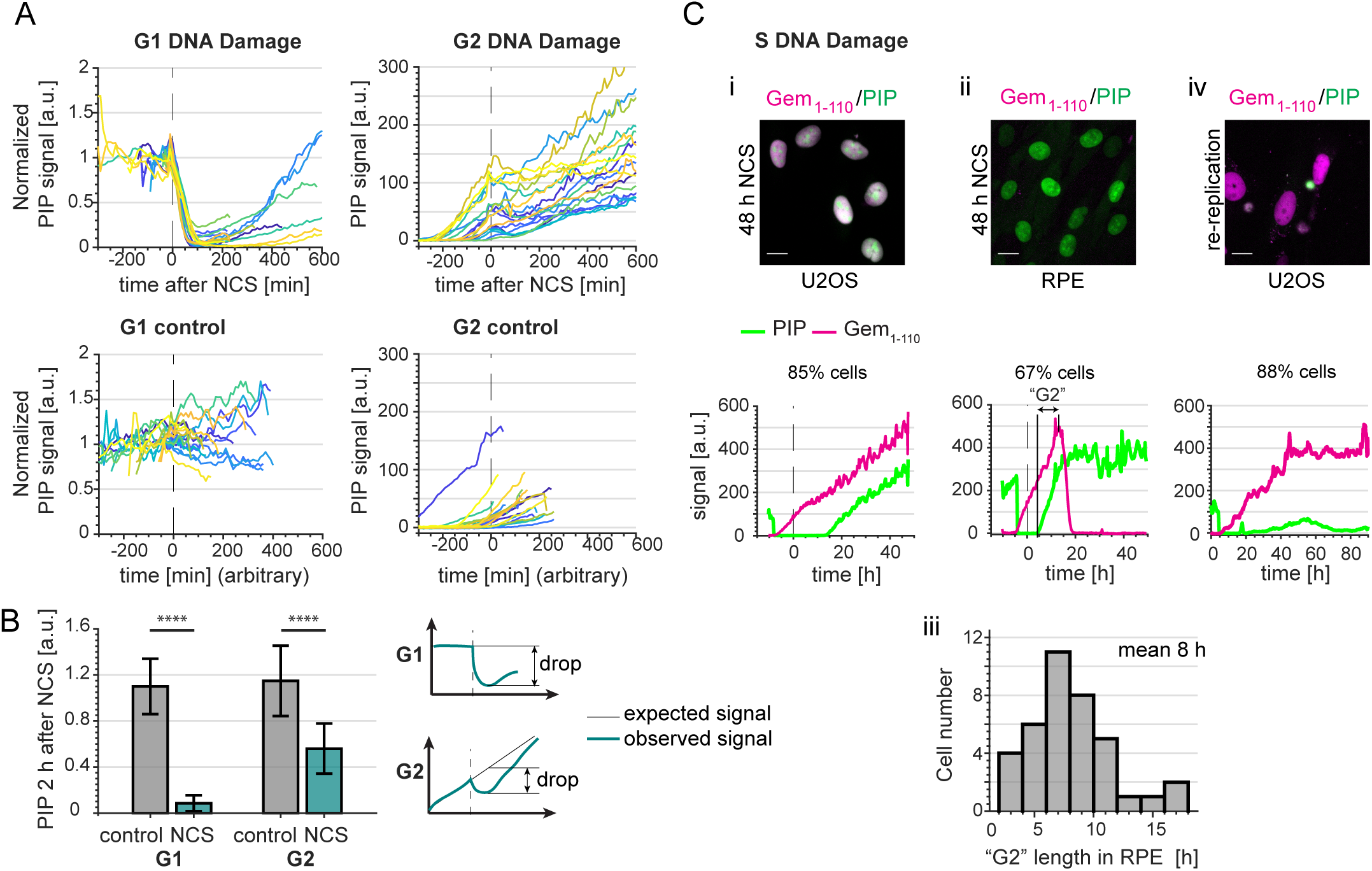
PIP-FUCCI dynamics after DNA damage. A. PIP-mVenus dynamics in control U2OS cells and cells treated with neocarzinostatin (NCS) (200 ng/ml). Upper graphs show PIP-mVenus traces from cells treated with NCS in G1 (left, n=19) or G2 (right, n=18). Lower graphs are traces from control cells in G1 (left, n=18) or G2 (right, n=22) phases. The traces end when cells enter S phase or mitosis respectively; x axis is experiment time in minutes, time of NCS addition is 0 (arbitrary time point was selected for control cells).
B. Quantification of PIP-mVenus decline 120 minutes after NCS treatment in U2OS cells in G1 and G2 phases. (Left) Bars are the ratios of detected versus expected PIP-mVenus levels assuming steady signal in G1 phase and linear increase in G2 phase. (Right) Cartoon showing the principle of this analysis. As a control traces from untreated U2OS cells were analyzed with a randomly selected point of mock treatment (for further details see Materials and Methods). Bar graphs show mean +/- standard deviation, Wilcoxon rank sum test, p<0.0001 (G1 control n=122, G1 NCS n=19, G2 control n=121, G2 NCS n=14).
C. Example micrographs and single cell quantification of PIP-FUCCI reporters in U2OS (i, n=48) and RPE-hTert (ii, n=59) cells 48 hrs after treatment with NCS. Dashed line indicates time of NCS addition x axis is real time in hours. (iii) histogram from NCS-treated RPE-hTert cells as in ii plotting hours between the end of S phase defined by PIP degradation and the start of degradation of mCherry-Gem_1-110_ in the absence of cell division (i.e. “mitosis skipping”) (iv) U2OS cells induced to re-replicate by overproducing full-length Cdt1 (n=32).

We next investigated the long-term PIP-FUCCI dynamics in response to NCS treatment in S phase. Because the PIP-mVenus signal was already below the detection threshold, we did not observe a further decrease after NCS treatment (e.g. Fig. 5Ci lower panel, first ~10 hrs after NCS). U2OS cells treated with NCS in S phase subsequently arrested and steadily accumulated both PIP-mVenus and mCherry-Gem_1-110_ until the end of the imaging (example trace shown in Fig. 5Ci). The rate of increase of both reporters was similar to untreated G2 cells. This behavior is similar to our previous analysis of the G2 arrest in S phase-damaged U2OS cells (18). RPE-hTert cells treated with NCS in S phase did not follow a similar pattern of sensor accumulation. After a variable arrest period with a mean of 8 hrs, cells expressed a stable amount of PIP-mVenus but then degraded mCherry-Gem_1-110_ (Fig. 5Cii and iii). The rapid mCherry-Gem_1-110_ degradation is consistent with APC^Cdh1^ activation despite the fact that the cells did not undergo nuclear envelope breakdown or cell division. We interpret this behavior as an indicator that cells could not maintain a full checkpoint arrest in G2 and reentered a G1 like state, bypassing mitosis and reactivating APC^Cdh1^ (42–44).

Finally, we examined the reporter dynamics in cells that had been transduced with a full-length, hyperactive Cdt1 to induce aberrant origin re-licensing after G1 phase. Origin re-licensing and re-firing is an endogenously-generated form of DNA damage that can occur in S or G2 (6, 45–51). We identified a subpopulation of cells that accumulated mCherry-Gem_1-110_, but without consistent accumulation of PIP-mVenus. For these cells, PIP-mVenus reporter fluorescence rose intermittently but did not accumulate as it did in G2 cells (Fig. 5Civ). This pattern is consistent with perpetual DNA replication and targeting of CRL4^Cdt2^ substrates (as we had previously reported), and cells remaining in an S-phase like state (49). Taken together, these examples show that the relative accumulation and dynamics of the PIP-FUCCI reporters can distinguish cell type-specific and DNA damage source-dependent arrest phenotypes.

### S phase onset can precede APC^Cdh1^ inactivation

Cells of different origins often have different cell cycle dynamics. This variation is particularly true for oncogenically transformed cells such as the osteosarcoma-derived U2OS cells, which have shorter G1 phase and longer S phase compared to untransformed RPE-hTert cells (Fig. 4C). In addition to affecting overall cell cycle phase lengths, cancer-associated genetic alterations or developmental signals could also affect temporal relationships of other events during the cell cycle. To explore this idea further, we introduced PIP-FUCCI into a third cell line, the glioblastoma multiforme-derived T98G line and compared the resulting dynamics to those of RPE-hTert and U2OS cells (52).

Interestingly, we observed variation in the relative timing between the onset of mCherry-Gem_1-110_ accumulation and PIP-mVenus degradation in each of the three cell lines (Fig. 6A and S3). Since PIP-mVenus degradation is a near-exact marker of S phase onset (Fig. 4C), the discrepancy means that mCherry-Gem_1-110_ is an imprecise marker of the G1 to S transition. In non-transformed RPE-hTert cells, mCherry-Gem_1-110_ was easily detectable before the degradation of PIP-mVenus whereas in U2OS cells it occurred at nearly the same time (Fig. 6A). In general, we detected mCherry-Gem_1-110_ rise earlier in cells with higher maximum reporter expression (Fig. S4). This correlation further emphasizes that, like the Cdt1-derived SCF reporters, Gem_1-110_-based reporters are not consistent markers of S phase entry *per se.* Surprisingly, mCherry-Gem_1-110_ detection was significantly delayed in comparison to PIP-mVenus in T98G cells (Fig. 6A and quantified in 6E). We refer to the difference between the onset of S phase (PIP_mid_) and in the initial rise of mCherry-Gem_1-110_ signal as the “Gem_1-110_ lag”, and its value can be positive (PIP degradation occurs first) or negative (Gem accumulation occurs first). The mCherry-Gem_1-110_ reporter is an APC^Cdh1^ substrate, and APC^Cdh1^ inactivation (i.e. mCherry-Gem_1-110_ accumulation) is thought to be required for S phase entry (53). Frequent and substantial Gem_1-110_ lag in T98G cells (Fig. 6E, left) suggests that APC^Cdh1^ inactivation does not always precede S phase entry.

**Figure 6.**
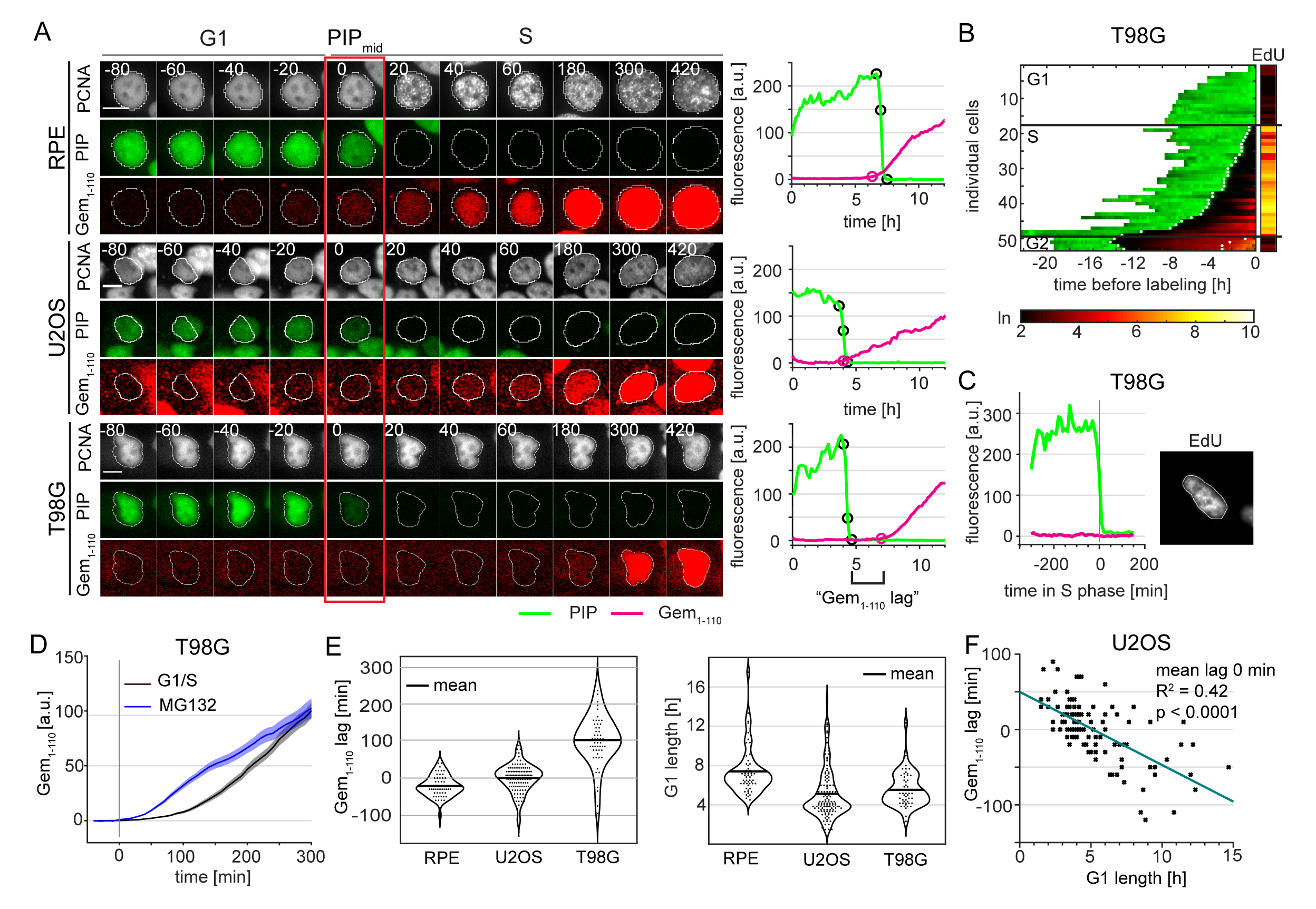
Variable APC^Cdh1^ reporter accumulation relative to the start of S phase in different cell lines. A. (Left) Micrographs of a single representative cell of RPE-hTert, U2OS, and T98G cells expressing PIP-FUCCI and mTurq2-PCNA; the red rectangle marks the frame of PIP_mid_. (Right) Quantification PIP-FUCCI reporter intensities the cells shown at left. PIP_max_, PIP_mid_, PIP_min_, and Gemrise are indicated by circles. Delay of Gemrise relative to both PIP_mid_ is defined as “Gem_1-110_ lag”, and is an indicator of the relative timing of APC^Cdh1^ inactivation and S phase onset.
B. Heat map of the signal intensity values of PIP-mVenus (green) and Geminin_1-110_ in asynchronously growing T98G cells before EdU labeling in the final 30 minutes. EdU intensity value (right, logarithmic scale) of the individual cells. White dots indicate the points of degradation and re-accumulation of PIP-mVenus.
C. PIP-FUCCI signals from a single T98G cell before EdU labeling in early S phase. Cells were labelled with EdU for the final 30 minutes of the experiment.
D. Comparison of mCherry-Gem_1-110_ accumulation at the G1/S transition (black, n=53) or after proteasome inhibition in G1 phase (MG132, 20 μM, blue, n=51) in T98G cells. Traces of asynchronous cells were aligned with time point zero corresponding to PIP_mid_ point for G1/S cells and the moment of drug exposure for MG132 treated cells. Traces show mean +/- SEM of the populations.
E. (Left) Quantification of the mean Gem_1-110_ lag in untreated, asynchronous RPE-hTert (n=56), U2OS (n=120), and T98G cells (n=51). Gem_rise_ that precedes S phase onset has a negative value. (Right) Distributions of G1 length in different cell lines as measured by PIP-mVenus dynamics (RPE-hTert n=56, U2OS n=120, T98G n=51).
F. Correlation between G1 length and Gem_1-110_ lag in untreated asynchronous U2OS cells (n=120).

We considered the possibility that the unexpected dynamics of the Gem_1-110_ APC^Cdh1^ reporter relative to S phase may be either an issue with S phase detection or a technical issue with mCherry-Gem1-110 detection itself. To determine if the S phase sensor PIP-mVenus was degraded prematurely in G1, we imaged asynchronous T98G cells expressing PIP-FUCCI for 24 hours before adding EdU for the last half hour of imaging. We fixed and stained for DNA synthesis (EdU) and DNA (DAPI). We matched the individual cells pre-fixing and post-fixing and confirmed that PIP was expressed throughout the entirety of G1 (Fig. 6B). We found no PIP-mVenus-negative cells that were not also EdU positive. More importantly, we found multiple examples of EdU positive cells that did not yet have detectable mCherry-Gem_1-110_ (Fig 6C and S3B). Thus, the mCherry-Gem_1-110_ reporter was indeed detected well after S phase onset in a substantial subpopulation of T98G cells.

We next tested the possibility that mCherry-Gem_1-110_ was indeed accumulating at S phase onset but that we could not detect it. Delayed detection could be exacerbated by the time needed for fluorescent protein maturation (54).To test the hypothesis that the delay in mCherry-Gem_1-110_ could be simply the long maturation time of mCherry or total reporter expression (Fig. S4), we treated asynchronous T98G cells with the proteasome inhibitor MG132 and quantified mCherry-Gem_1-110_ accumulation in cells that were in G1 phase (i.e. APC^Cdh1^ active) at the moment of treatment. We then compared the dynamics of accumulation of mCherry-Gem1-110 in cells with inhibited proteasome to the dynamics in early S. The mCherry-Gem_1-110_ accumulation was faster in MG132 treated cells than in cells entering S phase (Fig. 6D). The more rapid detection of mCherry-Gem_1-110_ in the MG132-treated cells indicates that the delayed detection in S phase T98G cells is not because we could not have detected the reporter. Since cells in both experiments were imaged using the same settings, the apparent “lag” in mCherry-Gem_1-110_ detection cannot be explained by low sensitivity or the time needed for mCherry maturation. The subset of lagging cells were indeed in S phase with (at least partially) active APC^Cdh1^.

We also detected a correlation between G1 length of individual U2OS cells and mCherry-Gem_1-110_ lag (Fig. 6F). Individual U2OS cells with short G1 lengths were more likely to begin mCherry-Gem_1-110_ accumulation after the beginning of S phase, whereas cells with longer G1 phases were more likely to accumulate mCherry-Gem_1-110_ coincident with (i.e. lag=0) or preceding (i.e. negative lag) S phase onset. Even though the average lag of the U2OS population was close to zero, we did identify cells with delayed mCherry-Gem_1-110_ accumulation, and these cells had the shortest G1 phases. The correlation between G1 duration and the lag in mCherry-Gem_1-110_ accumulation, prompted us to hypothesize that shortening G1 could increase the mCherry-Gem_1-110_ lag.

To test this idea, we analyzed the behavior of PIP-FUCCI in cells with significantly shortened G1 phase. We chose RPE-hTert for this experiment because they normally have a relatively long G1 phase (Fig. 4C and 6E) which can be significantly shortened upon overproduction of Cyclin E (2, 55, 56). For that reason, we expressed PIP-FUCCI in RPE-hTert cells that also bear a stably integrated doxycycline inducible Cyclin E cDNA and cultured the cells in doxycycline prior to and during imaging. Moderate Cyclin E overproduction dramatically shortened G1 phase (Fig. 7A, right). Most importantly, most of these cells accumulated mCherry-Gem_1-110_ well after the onset of DNA replication (Fig. 7B, C, S5A and Supplementary Movies 3 and 4). These cells showed both degradation of PIP-mVenus and incorporation of EdU prior to any detectable increase in mCherry-Gem_1-110_ (Fig. 7D). To again ensure that this was not due to inadequate mCherry-Gem_1-110_ detection in early S phase, we compared the lag times between control cells, cyclin E-overproducing cells, and cells treated with MG132 in G1 (Fig. 7E). We set the onset of DNA replication and the addition of MG132 arbitrarily at time point zero. As before (Fig. 6D), the control cells accumulated mCherry-Gem_1-110_ before the onset of S phase (black traces). Cells treated with MG132 accumulated mCherry-Gem_1-110_ earlier than cells overexpressing cyclin E.

**Figure 7.**
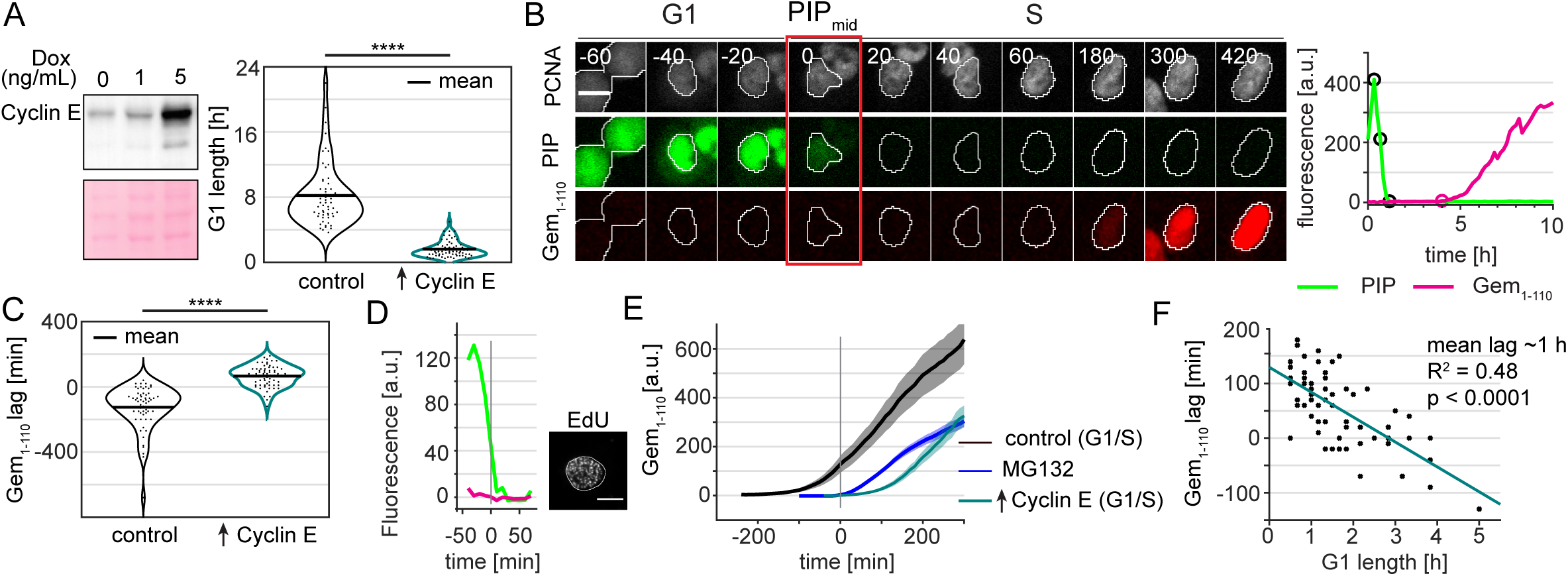
Delayed APC^Cdh1^ reporter accumulation relative to the start of S phase in RPE cells overproducing cyclin E. A. (Left) Cyclin E overproduction in RPE cells. Clonally-selected RPE-hTert cells with a stably integrated, doxycycline-inducible cyclin E cDNA were treated with 1 ng/mL, 5 ng/mL doxycycline or vehicle control for 48 hours. Whole cell lysates were immunoblotted for Cyclin E or stained for total protein with Ponceau S. (Right) Distributions of G1 lengths in control RPE-hTert (n=58) and in their isogenic counterparts overproducing Cyclin E (n=65).
B. (Left) Example cell showing Gem_1-110_ lag in PIP-FUCCI RPE cells stably transduced with a doxycycline-inducible cyclin E cDNA expression construct and cultured in 5 ng/ml doxycycline. Numbers indicate minutes from the G1/S transition as measured by PIP-mVenus dynamics; scale bar 10 μm. (Right) Quantification of PIP-FUCCI reporter intensities in the cell shown at left. PIP_max_, PIP_mid_, PIP_min_, and Gem_rise_ are indicated by circles.
C. Distributions of mCherry-gem_1-110_ lag in control or cyclin E-overproducing cells. Positive lag indicates mCherry-gem_1-110_ accumulation started after S phase. Kolmogorov-Smirnov test, ^⋆⋆⋆⋆^ p<0.0001.
D. PIP-FUCCI signals from a single RPE Cyclin E-overproducing cell before EdU labeling in early S phase.
E. Comparison of mCherry-Gem_1-110_ accumulation at the G1/S transition in RPE control (black, n=31) and RPE cells overproducing Cyclin E (teal, n=32) relative to with RPE cells treated with MG132 in G1 phase (20μM MG132, blue, n=59). Traces of asynchronous cells were aligned with time point zero corresponding to PIP_mid_ point for G1/S cells and the moment of drug exposure for MG132 treated cells. Traces show mean +/- SEM of the populations.
F. Correlation between G1 length and Gem_1-110_ lag in RPE cells expressing cyclin E (n=65).

These results indicate that the lag in mCherry-Gem_1-110_ accumulation in RPE cells overproducing Cyclin E is due to a significant remainder of APC^Cdh1^ activity during early S phase. Moreover, the individual cells with the shortest G1 phase lengths had the largest mCherry-Gem_1-110_ lag. Interestingly, these cells also had prolonged S phase suggesting that APC^Cdh1^ activity that extends past the G1/S transition may cause perturbed replication dynamics in S phase (Fig. 7F and S5B, C). We also shortened G1 phase by and induced mCherry-Gem_1-110_ lag in U2OS cells by depleting the p53 tumor suppressor (Fig. S6). The stronger effect on G1 length from cyclin E overproduction compared to p53 depletion corresponded to a greater magnitude of induced lag in mCherry-Gem_1-110_ accumulation (Fig. 7C and S6B). From these experiments, we conclude that the relative timing of APC^Cdh1^ inactivation and the start of S phase is variable by cell line and among individual cells. This timing is subject to genetic perturbation, and the decoupling of APC^Cdh1^ inactivation from DNA replication onset may lead to a less robust S phase. We further note that these observations would be difficult to discern without the ability to precisely mark the relevant cell cycle transition events in live, single cells.

## Discussion

In this report we make three principal contributions: 1) We provide a detailed mechanistic analysis of the widely-used FUCCI reporters that are based on human Cdt1_30-120_ fusions, 2) We validate a new replication-coupled destruction reporter and compare it to another replication-based PCNA reporter, and 3) Using these new tools, we demonstrate frequent and genetically inducible decoupling of the timing of APC^Cdh1^ inactivation relative to the G1/S transition. We describe here some of the implications of these results and suggest strategies for selecting and interpreting mammalian cell cycle reporters.

### Interpreting original FUCCI dynamics

The FUCCI Cdt1_30-120_ fusion is a reporter of SCF^Skp2^ activity *per se,* but does not precisely define the boundaries of S phase (Table 1). The fusion continues to accumulate after S phase starts, and its abundance is generally low from late S through G2 and M phase (Fig. 1E). This pattern reflects the abundance and activity of the Skp2 substrate adapter protein, which itself is a substrate of APC^Cdh1^ (57, 58). In that sense, Cdt1_30-120_ is in fact an indirect reporter of APC^Cdh1^. This behavior precludes using Cdt1_30-120_ to mark either of the S phase boundaries. Moreover, the peak of Cdt1_30-120_ relative to S phase time varies with the maximum level of reporter expression, so it is not possible to correctly mark S phase by subtracting or adding a constant time from the peak. In other words, investigators can assume that S phase started some time before the Cdt1_30-120_ peak, but they cannot reliably determine the magnitude of this offset in individual cells, even in clonally-selected populations. This behavior may be problematic for studies in which distinguishing G1 cells from early S phase cells is important.

**Table 1.** Cell cycle reporters for use in mammalian cells.

| Reporter | Pattern |  | Advantages | Considerations |
| --- | --- | --- | --- | --- |
| **PCNA**<br>(or PCNA nanobody (14)) | G1 | diffuse nuclear | • DNA replication sensor<br>• Single fluorescence channel for all phases (time lapse only)<br>• high-contrast S/G2 transition marker | • requires high resolution images<br>• requires regular nuclear morphology and monolayer culture<br>• subtle G1/S change |
|  | S | punctate |  |  |
|  | G2 | diffuse nuclear |  |  |
|  | M | diffuse whole cell |  |  |
| **Cdt1<sub>30-120</sub>**<br>(or Cdt1<sub>16-120</sub>) | G1 | increasing | • SCF<sup>Skp2</sup> activity sensor | • requires T or E in linker (at 29)<br>• slow degradation in S phase<br>• very sensitive to reporter expression levels |
|  | S | peaks after G1/S |  |  |
|  | G2 | low |  |  |
|  | M | low |  |  |
| **PIP** | G1 | stable | • DNA replication sensor<br>• single fluorescence channel for all phases (time lapse only)<br>• high-contrast G1/S marker<br>• insensitive to reporter expression levels | • absent in S phase (requires 2<sup>nd</sup> marker for automated tracking)<br>• sensitive to high levels of exogenous DNA damage in G1 |
|  | S | absent |  |  |
|  | G2 | increasing |  |  |
|  | M | diffuse whole cell |  |  |
| **Geminin<sub>1-110</sub>**<br>(or Geminin<sub>1-60</sub>) | G1 | absent | • APC activity sensor<br>• distinguishes S and G2 from G1<br>• Geminin<sub>1-60</sub> labels cytoplasm (cell shape) | • unreliable marker of S phase entry<br>• somewhat sensitive to reporter expression levels |
|  | S | increasing |  |  |
|  | G2 | increasing |  |  |
|  | M | decreasing |  |  |

It is important to note that FUCCI strategies for use in *Drosophila melanogaster* (“Fly-FUCCI”) and *Danio rerio* (“zFUCCI”) both contain PIP degrons from dmE2F and zCdt1 respectively (9–11). These derivatives are subject to replication-coupled destruction similar to the PIP-mVenus reporter here as well as two recent “hCdt1(1/100)” reporters from Sakaue-Sawano 2017. These PIP degron-containing reporters are *bona fide* S phase reporters. A Cdt1-based cell cycle reporter for use in plants has also been described using a C-terminal fragment of plant Cdt1. This fragment accumulates in S and G2 similar to human Gem_1-110_, has no readily-identifiable PIP degron, and contains motifs that suggest plant Cdt1 may be an APC substrate rather than a CRL4^Cdt2^ substrate (59).

We also demonstrate here that oscillations of the original FUCCI Cdt1_30-120_ reporter are entirely dependent on a non-native glutamate residue corresponding to position T29 in the human Cdt1 phosphodegron. A reporter lacking the glutamate but retaining the potentially redundant S31 failed to oscillate (Fig. 2C), and the inability of S31 to substitute for T29 is consistent with a prior study (26). We presume that the glutamate is a serendipitous contribution from the vector sequence in the original expression construct. We postulate that the reason the original CdtW1_30-120_ reporter only oscillated when fused to one of the four tested fluorescent proteins is that other expression vectors lacked the phosphomimetic residue. Failure of reporters based on human Cdt1_30-120_ to oscillate in zebrafish or fruit flies may also be related to the vector sequences used in those attempts and not an intrinsic property of either human Cdt1 or the suggested species-specific differences in SCF^Skp2^ (9, 10). Interestingly, glutamate and threonine at position 29 were equivalent for generating a SCF^Skp2^ reporter. In native Cdt1, T29 is phosphorylated by Cyclin A/Cdk2 during S phase, and the phosphorylation creates a binding site for the Skp2 substrate targeting subunit of SCF (26). It was possible that our test reporters with threonine instead of glutamate in that position would have different dynamics since the threonine-bearing reporters integrate both cyclin A-mediated phosphorylation and SCF^Skp2^ activity whereas the phosphomimetic FUCCI requires only Skp2 activity. The patterns of the FUCCI reporter and both phospho-dependent SCF reporters (Cdt1_16-120_ and FUCCI_E29T_) were remarkably similar however (Fig. 1E, 2B, and 2D). We suggest therefore that the threonine-bearing reporters are likely fully phosphorylated, at least in the U2OS cells we analyzed. By extension, cyclin A/Cdk2 activity is delayed from the onset of S phase and is not rate-limiting for targeting by SCF^Skp2^; this relationship may also apply to endogenous Skp2 targets.

### PIP-FUCCI is a precise marker of S phase

Three recent studies used fluorescent fusions to PCNA or a PCNA nanobody to track the formation of DNA replication foci, and thus S phase, in living cells (14, 15, 60). These approaches take advantage of the well-established patterns of DNA replication “factories” in mammalian cells that are readily detected as foci of nucleotide incorporation or by probing for replication fork proteins such as PCNA (13, 17, 19, 61). Changes in PCNA localization are sufficient to mark the boundaries of S phase, and successful use of this strategy can precisely mark all four major cell cycle transitions using a single fluorescence channel. One challenge for analyzing cell cycle progression by PCNA localization patterns is devising an image analysis pipeline that is unbiased, quantitative, and efficient. Burgess *et al.* quantified the number of PCNA foci within S phase, but this approach requires high magnification and high resolution images. We describe here the use of total PCNA signal variance which is amenable to lower-magnification/lower light-intensity images.

Despite the powerful advantages of using a single localization-based fusion to mark all cell cycle transitions, the requirements for this approach are still beyond what is feasible in some studies. We thus sought to create and thoroughly validate a simple intensity-based cell cycle reporter strategy. The G1/S transition and S/G2 transition reporter in our PIP-FUCCI system leverages a well-characterized replication-coupled destruction pathway in metazoan cells for S phase degradation via CRL4^Cdt2^. Degradation of the reporter relies solely on interaction of the human Cdt1 PIP degron with PCNA when PCNA is loaded onto DNA. Reporter dynamics are thus a direct physical readout of DNA replication *per se.* Moreover, the very rapid degradation conferred by the Cdt1 PIP degron should make the G1/S transition particularly easy to identify in low-resolution images or even in tissues or organoids. Of particular importance, PIP-fusion reporter dynamics are unaffected by reporter expression within a broad range. In all analyzed cells where the reporter was expressed above background detection levels, it marked both the G1/S and S/G2 transitions as accurately as PCNA variance did. This robustness is in stark contrast to each of the SCF^Skp2^ reporters that had very different dynamics in different cells depending on the level of reporter expression. The PIP degron reporter can mark the same transitions as PCNA in time-lapse images, although it does not distinguish early from late S phase. In addition, PCNA is a more robust mark of the S/G2 transition in cells with lower PIP reporter fluorescence intensity, since the fusion must rise above background to identify the transition to G2.

The near-perfect correlation of PIP-FUCCI dynamics with S phase means that the PIP degron reporter is insensitive to typical levels of endogenously generated DNA damage. We could trigger degradation of the PIP-mVenus reporter with relatively high doses of exogenous genotoxin however. Thus, in otherwise proliferating cells, PIP-degron degradation can be interpreted as a mark of S phase onset with confidence. Furthermore, even high doses of NCS could not induce degradation of the PIP degron reporter in G2 cells. This stability is expected given reports by us and others that the CRL4^Cdt2^ E3 ligase is inactivated by CDK1 during G2 and M phase (40, 41). Thus, like endogenous Cdt1, the PIP degron reporter accumulates in G2 phase both because PCNA is unloaded from DNA and because CRL4^Cdt2^ is inactive. We emphasize, however that the PIP degron reporter is not an accurate biosensor for *endogenous* full-length human Cdt1 dynamics. The SCF^Skp2^-based phosphodegron likely contributes very little to the normally rapid Cdt1 degradation at G1/S, given the dynamics of the SCF^Skp2^-based reporters described here. On the other hand, SCF^Skp2^ targeting and cyclin/Cdk binding may influence endogenous Cdt1 stability in late S phase and G2. Presumably, this activity is why the SCF^Skp2^ reporters all remain low in G2 phase, but full-length Cdt1 re-accumulates in many cell types (2, 62). In addition, phosphorylation at sites in the C-terminal third of full-length Cdt1 and binding to full-length endogenous geminin can stabilize Cdt1, at least in some cell lines and settings (62–65).

### Selecting cell cycle reporters

For studies requiring the identification of cell cycle phases in mammalian cells, we provide suggestions for selecting the appropriate fluorescent reporters (Fig. 8 and Table 1). Identifying cell cycle phases in <u>fixed</u> cells requires at least two fluorophores. We recommend either PIP-FUCCI described here or “FUCCI(CA)” from Sakaue-Sawano 2017. Both of these reporter pairs include the Geminin_1-110_ fragment which is necessary to distinguish G2 cells from G1 cells (e.g. Fig. 3F). The PIP-mVenus fusion contains 17 amino acids from Cdt1 which, to our knowledge, only binds PCNA and Cdt2, and it can be overproduced quite highly to distinguish S from G2 cells without any effect on either cell proliferation or the ability to precisely mark cell cycle transitions. FUCCI (SCA) is human Cdt1 1-100 which contains the strong cyclin A binding motif at amino acids 68-70. This inclusion raises the possibility of the reporter competing with endogenous cyclin A binding partners. The reporter also contains phosphorylation sites of unknown consequence and most of a negative regulatory domain of unknown mechanism that may influence Cdt1 chromatin binding (27, 66). The derivative reporter in the FUCCI(CA) pair, “hCdt1(1-100)Cy(-)”, has fewer of the considerations related to cycling binding, but the minimal PIP degron described here provides particularly robust identification of S phase transitions with no additional known interactions. Including a third reporter ensures that cells are visible even during early S phase when both reporters are quite low. For this third maker, we recommend PCNA or a fusion to histone H2B which mark the subphases of S phase and mitosis respectively.

**Figure 8.**
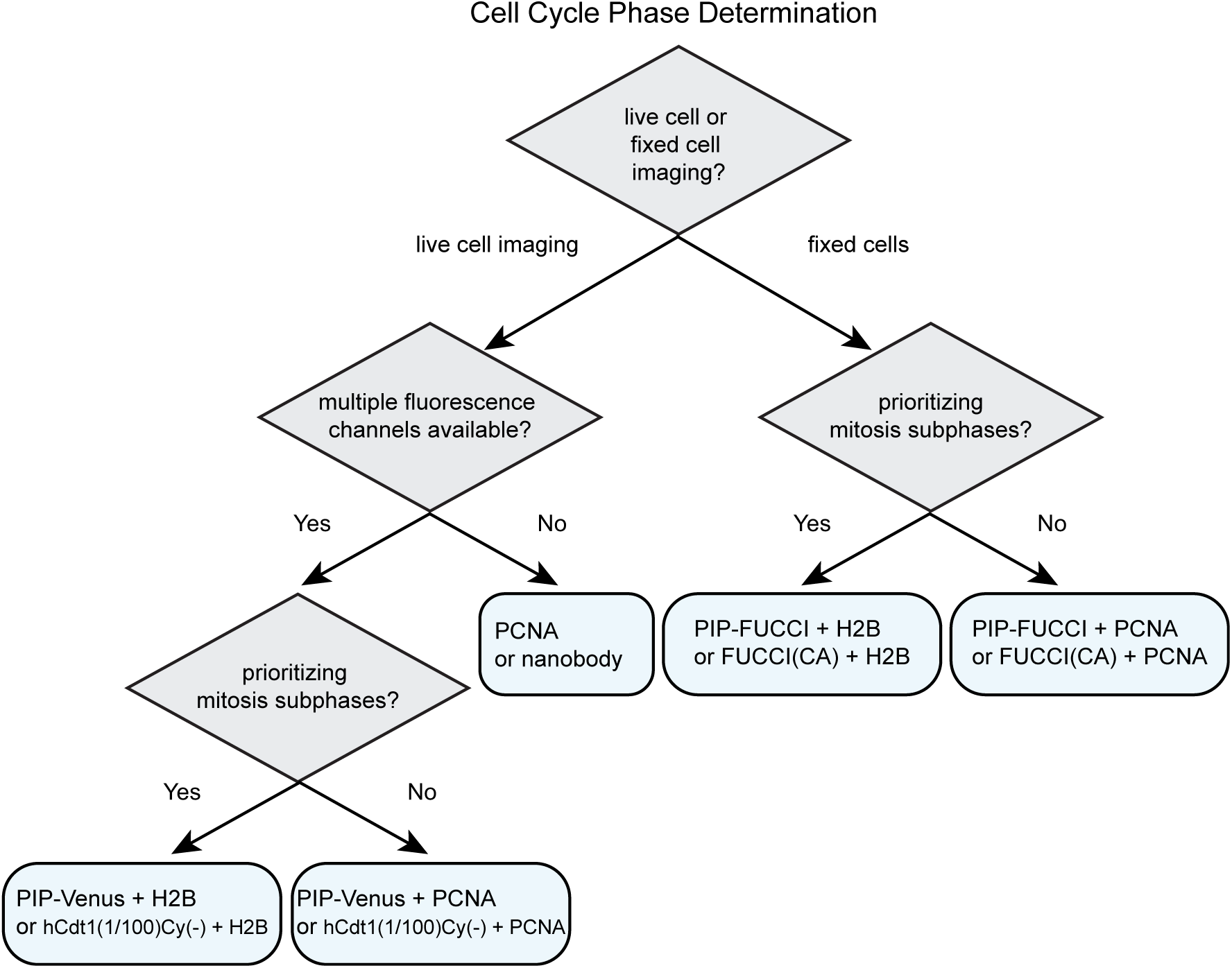
An illustrated strategy for selecting cell cycle phase reporters for use in mammalian cells. See Table 1 and Discussion for details.

For time-lapse microscopy, several options are now available for dynamic cell cycle phase analysis. If only one fluorescence channel is available, then a PCNA fusion or the PCNA nanobody or alternatively, the PIP degron fusion or hCdt1(1-100)Cy(-) can mark all four cell cycle phases; PCNA localization provides additional information about progression within S phase and does not require an additional channel for cell tracking. Relying on a single cell cycle reporter makes other fluorescence channels available for simultaneous analysis of additional reporters of interest, such as reporters of kinase activity (e.g. CDK), transcription (e.g. E2F), or ubiquitin ligase activity (e.g. SCF^Skp2^ or APC^Cdh1^) (4, 67, 68). Cell cycle phase analysis by PCNA localization requires time-lapse microscopy of sufficient resolution in cells with regular nuclear morphology to robustly detect the smaller PCNA foci in early S phase, but the dramatic dissolution of large foci at the end of S phase makes it an excellent marker of the S/G2 transition. When multiple fluorescence channels are available, we recommend including both a cell cycle phase reporter and a constitutively stable reporter for automated tracking (PCNA or histone H2B). Both the PIP degron reporter (i.e. PIP-mVenus) and Cdt1(1/100)Cy(-) are accurate reporters of the beginning and end of S phase, and the end of S phase is best visualized in cells with high reporter expression since it requires accumulation above the detection threshold.

### De-coupled events at the G1/S transition

APC^Cdh1^ is inactive in S phase, so it is not surprising that we find that the mCherry-Gem_1-110_ APC^Cdh1^ substrate reporter begins accumulating at or near S phase onset in most cells (53, 69–71). The accumulation of nearly all APC^Cdh1^ substrates during S phase and many previous studies finding that artificially depleting Cdh1 shortens G1 and accelerates S phase onset have all suggested that APC^Cdh1^ inactivation as a pre-requisite for S phase entry (53, 70, 72). Based on the prior assertion that APC^Cdh1^ inactivation and DNA replication initiation are mechanistically coupled, we were thus surprised to find that the correlation between the timing of mCherry-Gem_1-110_ accumulation and S phase entry can be quite variable, even in clonally-selected populations. The precise timing of mCherry-Gem_1-110_ accumulation is affected by the expression level of the reporter; cells that ultimately have high maximal mCherry-Gem_1-110_ levels cross the detection threshold earlier on average than cells with lower expression (Fig. S4). That detection threshold is a combination of the microscopy settings and the folding time for mCherry (~40 min). This technical consideration accentuates the utility of reporters that are degraded or relocalized (e.g. PIP-mVenus, or PCNA) rather than reporters that accumulate for precisely marking cell cycle transitions.

We postulate however that mCherry-Gem_1-110_ reporter detection limitations cannot account for all of the variability we detected. First, the accumulation of the PIP-mVenus fusion in G2 has similar detection considerations as those that apply to mCherry-Gem_1-110_, and we found no analogous lag between the end of S phase (as marked by PCNA) and the accumulation of PIP-mVenus (Fig. S6F). Second, we could genetically exacerbate the difference between S phase onset (measured by both PCNA foci and PIP-mVenus degradation) and APC/C reporter accumulation by overproducing cyclin E or depleting p53. Both of these biological manipulations have the effect of activating Cdk2; one direct (cyclin E) and the other indirect because p53-depleted cells have low levels of the Cdk inhibitor p21 (Fig. S6D). Third, even in otherwise unperturbed cells, we found a strong correlation between G1 length and the time of initial mCherry-Gem_1-110_ detection. We observed the same relationship with the Citrine-Gem_1-110_ reporter (unpublished observations). Thus even within a clonal population, cells with shorter G1 phases inactivated APC^Cdh1^ later relative to S phase onset. One interpretation of this correlation is that the pathway for APC^Cdh1^ inactivation is coordinated with, but not in lockstep with, the series of steps leading to DNA replication itself.

We cannot rule out the possibility of partial APC/C inactivation in those cells with significant Gem_1-110_ lag. Cells that started S phase with mCherry-Gem_1-110_ levels still quite low might have some intermediate APC^Cdh1^ activity that allowed partial accumulation of invisible critical APC^Cdh1^ substrates. One or more of these substrates may indeed be a strict pre-requisite for S phase onset, but could be required only at very low levels. Yuan et al. reported that Cdh1 depletion not only induced premature S phase entry, but the S phase that ensued was longer than normal with perturbed replication fork dynamics. We speculate that when APC^Cdh1^ inactivation precedes S phase entry cells execute a robust switch from G1 into S phase. Slow or delayed APC^Cdh1^ inactivation may result in a gradual and inefficient S phase entry that could promote genome instability. In support of the importance of the normal coupling of APC^Cdh1^ inactivation and S phase entry, cells with short G1 phases were more likely to inactivate APC^Cdh1^ after S phase began (Fig. 6F and 7F) but then spent more overall time in S phase (Supplementary Fig. S5C).

Because DNA replication is an essentially irreversible step, S phase entry and progression is tightly regulated. Detecting the G1/S transition in living cells is not a simple technological task however. PIP-FUCCI is a precise method for demarcating cell cycle phase transitions in mammalian cells. Relative to first-generation FUCCI reporters, the simplicity of the new PIP-mVenus fragment reduces the number of confounding factors that must be considered when designing and interpreting experiments. PIP-FUCCI reagents are available to researchers for straightforward cell cycle phase determination. We anticipate that the ability to clearly distinguish G1 from S phase and S phase from G2 in both fixed and live cells will be particularly useful for investigations focused on cell proliferation commitment and the maintenance of genome stability.

## Materials and Methods

### Cell culture and Cell line construction

HEK293T, T98G, and RPE1-hTert (RPE) cells were obtained from the American Type Culture Collection (ATCC). U2OS cells were a gift from M. Whitfield. All cells were passaged in Dulbecco’s modified Eagle’s medium (DMEM, Sigma) supplemented with 10% fetal bovine serum (FBS, Sigma), 1x Pen/Strep, and 4mM L-glutamine and incubated at 37°C in 5% CO_2_. U2OS and RPE-hTert lines were authenticated by STR profiling (ATCC) and all lines were confirmed to be mycoplasma free. Cell lines were selected using the following concentrations of antibiotics: puromycin-1 μg/mL for U2OS and T98G cells, 5-10 μg/mL for RPE cells; neomycin 100 μg/mL for RPE cells, 500 μg/mL for U2OS and T98G cells; hygromycin 150 μg/mL for U2OS and T98G cells; zeocin 100 μg/mL for U2OS and RPE cells, and blasticidin 10 μg/mL for U2OS, RPE, and T98G cells.

### Cell transfection

X-trmeGENE 9 was used to transfect ES-FUCCI into RPE cells using the manufacturers recommended protocol. X-tremeGENE HP was used to transfect ES-FUCCI into U2OS cells following manufacturers recommended protocol. Cells with stable integration of the plasmid were selected using 500 μg/mL G418 (Gibco). When necessary, colonies from single cells were selected by visual inspection for even expression of fluorescently tagged proteins.

### Cell transduction

To make stable cell lines using viral vectors initially HEK293T cells were transfected with pVSVG, pΔNRF (Gifts from Dr. J. Bear), and the viral vector of interest, using 50 μg/mL Polyethylenimine-Max (Aldrich Chemistry). Viral supernatant was transduced with 8 μg/mL Polybrene (Millipore) into U2OS cells for four hours or into RPE1 or T98G cells overnight.

### Plasmid construction

All constructs were made using either the Gateway cloning method (Invitrogen) or by Gibson Assembly (New England Biolabs, NEB) following standard protocols. PCR fragments were amplified using Q5 polymerase (NEB). DNA fragments were isolated using the Qiaprep spin miniprep kit (Qiagen). Plasmids were transformed into either DH5α or Stbl2. pENTR constructs were combined with the expression constructs: pLenti PGK Hygro DEST (w530-1), pLenti CMV Blast DEST (706-1), pLenti PGK Puro DEST (w529-2), and pLenti PGK Neo DEST (w531-1) were gifts from Eric Campeau and Paul Kaufman (Addgene plasmids # 19066, 17451, 19068, and 19067 respectively) using LR Clonase (Invitrogen). pInducer20-Cyclin E1 was described previously (73). Plasmids were validated via sequencing (Eton Biosciences) for the desired insert using appropriate primers. The cyclin E cDNA expression construct has been described previously (73).

### Total lysate

Total lysate was collected via trypsinization. Cells were washed with 1X phosphate buffer solution (PBS) and then centrifuged at 1700 × g. For total protein lysates, cells were lysed on ice in CSK buffer (300 mM sucrose, 100 mM NaCl, 3 mM MgCl2, 10 mM PIPES pH 7.0) with 0.5% triton X-100 and protease and phosphatase inhibitors (0.1 mM AEBSF, 1 mg/ mL pepstatin A, 1 mg/ mL leupeptin, 1 mg/ mL aprotinin, 10 mg/ ml phosvitin, 1 mM b-glycerol phosphate, 1 mM Na-orthovanadate). Cells were centrifuged at 13,000 × g at 4°C, then the supernatants were transferred to a new tube for a Bradford Assay (Biorad) using a BSA standard curve. Alternatively, an approximately equal number of cells were collected using SDS loading buffer.

### Immunoblotting

Immunoblotting samples were diluted with SDS loading buffer (final: 1% SDS, 2.5% 2-mercaptoethanol, 0.1% bromophenol blue, 50 mM Tris pH 6.8, 10% glycerol) and boiled for 5 min. Samples were run on SDS-PAGE gels, then the proteins transferred onto polyvinylidene difluoride membranes (PVDF) (Thermo-Fisher). Membranes were blocked at room temperature for 1 hr in either 5% milk or 5% BSA in Tris-Buffered-Saline-0.1%-tween-20 (TBST). After blocking, membranes were incubated in primary antibody overnight at 4°C in either 1.25% milk or 5% BSA in 1X TBST. Blots were washed with 1X TBST then incubated in HRP-conjugated secondary antibody in either 2.5% milk or 5% BSA in 1X TBST, washed with 1X TBST, and then membranes were incubated with ECL Prime (Amersham) and exposed using Chemi-Doc (Bio-Rad). Equal protein loading was verified by Ponceau S staining (Sigma-Aldrich). Antibodies used for immunoblotting were: Cyclin E1 (Cell Signalling Technologies, Cat# sc-198), p53 (Santa Cruz Biotechnologies, SC-126), p21 (Santa Cruz Biotechnologies, SC-397-G), and PCNA (Santa Cruz Biotechnologies, SC-25250).

### Live Cell imaging

Prior to imaging cells were plated on glass-bottom plates (Cellvis) #1.5 in FluoroBrite DMEM (Invitrogen) supplemented with FBS, L-glutamine, and penicillin/streptomycin (imaging media). Imaging was performed using a Nikon Ti Eclipse inverted microscope using Plan Apochromat dry objective lenses 20x (NA 0.75) or 40x (NA 0.95) and the Nikon Perfect Focus System. Images were captured using an Andor Zyla 4.2 sCMOS detector with 12 bit resolution. Cells were imagined in a humidified chamber (Okolabs) at 37°C with 5% CO_2_. All filter sets were from Chroma, CFP - 436/20 nm; 455 nm; 480/40 nm (excitation; beam splitter; emission filter), YFP - 500/20 nm; 515 nm; 535/30 nm; and mCherry - 560/40 nm; 585 nm; 630/75 nm. Images were collected either every 6 or 10 minutes using NIS-Elements AR software. No photobleaching or phototoxicity was observed in cells imaged with this protocol.

In proteasome inhibition experiments asynchronous populations of cells were treated with 20μM MG132 (Sigma). To study mCherry-Geminin_1-110_ increase, cells in G1 phase at the moment of treatment were selected for analysis.

### EdU labeling

Cells were plated on glass-bottom plates 24 hours prior to imaging in imaging media. After ~20 hours the media was changed and the cells were allowed to acclimate to the fresh media for ~4 hours. Cells were imaged using standard conditions for 22 hours (U2OS, T98G) for cell cycle progression analysis or treated with Dox and imaged for 5 hours (RPE-hTert Cyclin E). At the end of the movie (T98G, RPE Cyclin E) or after completion of the movie (U2OS) the culture media was supplemented with 10 μM EdU (Santa Cruz Biotechnology) for 30 minutes. After the EdU incubation, cells were washed with PBS and fixed with 4% paraformaldehyde (Electron Microscopy Sciences) for 15 minutes at room temperature. Cells were washed twice with 1% BSA-PBS before permeabilization with 0.5% Triton x-100 (Sigma) in 1% BSA-PBS for 15 minutes at room temperature. Cells were then washed twice with 1% BSA-PBS before a 30 minute incubation in PBS containing 1 mM CuSO_4_, 1 μM Alexa647-azide, and 100 mM ascorbic acid. Cells were then washed with 1% BSA + 0,5% Triton X-100 before an hour incubation in 1% BSA + 0.5% triton x-100 with 100 μg/mL RNase and 1 μg/mL DAPI (Life Technologies). After fixing and staining the cells were reimaged for EdU incorporation and DAPI staining at the same well locations as in the live cell movie so that the same cells could be used for the movie and fixed imaging analysis. Cells were scored for EdU incorporation vs cell cycle position in the live cell movie.

### siRNA

Cells were plated one day prior to imaging in FluoroBrite DMEM with FBS and L-glutamine at a density of 25,000 cells per well of a 6-well dish. The next morning Dharmafect 1 (Dharmacon) was diluted in FluoroBrite DMEM with the appropriate siRNA according to the manufacturer’s instructions. After changing the media on each well, diluted Dharmafect 1/ FluoroBrite / siRNA mixtures were added dropwise. The final concentration for each siRNA was 20 nM. Cells were imaged approximately 4 hours post siRNA addition with no additional media changes.

### DNA damage

Cells were treated with Neocarzinostatin (NCS, Sigma N9162) at final concentration of 200 ng/ml (U2OS) or 100 ng/ml (RPE-hTert). To quantify the relative loss of PIP-mVenus signal due to DNA damage, we calculated the ratio of signal intensity 120 minutes after NCS addition to the expected signal intensity calculated by extrapolating the PIP-mVenus signal before treatment. For cells in G1 the expected signal levels in untreated cells were calculated by assuming no change over time. The expected signals for untreated G2 cells were calculated assuming a linear increase in signal intensity over time.

### Image Analysis

Image analysis was done using Fiji (version 1.51n, ImageJ NIH) software (74) and Matlab (R2017b, MathWorks).

### Tracking and Segmentation

Individual cells were segmented and tracked in time-lapse movies by a user-assisted approach. Prior to analysis, all movies were pre-processed using rolling ball background subtraction. Background subtracted movies were kept in a 16-bit format, and residual background values resulting from noise clipping were subtracted in a separate step. Individual tracks were collected using a set of in-house developed ImageJ scripts that facilitate manual tracking by hovering with a mouse cursor over a selected cell while the movie advances at a chosen speed. Using user-defined tracks, nuclear regions of interest (ROI) were segmented automatically based on intensity of selected tracking channel (PCNA) followed by separation of touching nuclei by a watershed algorithm and a set of morphological operations to define oval shaped nuclear ROIs. In case of failed segmentation, the user could choose to provide manually defined polygons as replacement ROIs. The same set of ROIs was used to analyze all fluorescence channels.

### PCNA Analysis

The PCNA pattern was analyzed within nuclear ROIs as defined by tracking and segmentation (above). Images were smoothed using a Gaussian Blur filter (sigma=1x), darker regions (nucleoli) were eliminated using Remove Outliers algorithm (ImageJ) and PCNA patterns were enhanced using variance filter (sigma=2x). To avoid edge artifacts, intensity and standard deviation of variance image were measured within 70% of the central area of nuclear ROIs. A sum of mean and standard deviation of variance image (PCNA variance trace) showed the highest contrast at the beginning and the end of the S phase and was therefore used for cell cycle phase delineation. The beginning and end of S phase were detected automatically using the PCNA variance trace by smoothing this signal with filters of three different kernel sizes and looking for points that consistently showed highest change in value between G1/G2 and S phase.

### Phase warping of traces

The beginning and end of S phase were detected automatically based on PCNA variance trace as described above. Based on these values, the median durations of G1, S and G2/M phases were calculated for each analyzed cell population. These median values were used to calculate the percentage of total cell cycle time for each phase and create a cell cycle ‘blueprint’ of the population. Traces of individual cells were divided into G1, S, and G2/M, interpolated linearly within those given phases over the number of time points defined by the corresponding blueprint and then assembled to create a single phase warped signal.

### PIP-FUCCI trace annotation

Important points within PIP and Geminin_1-110_ signals (min, mid, max and rise) were detected automatically in each original trace (without phase warping). The signals were normalized to the maximum observed within a cell cycle. Alternatively, in experiments in which full cycle traces were not available max signal for PIP-mVenus was defined as maximum value observed in G1 phase and for mCherry-Geminin_1-110_ an arbitrary point of normalization was set at 5 h after the studied event (drug treatment or phase transition as defined by PIP-mVenus). Signals were smoothed with a kernel size 3 before annotation. ‘PIP min’ was defined as the first point within the signal that drops below a 5% threshold while ‘PIP rise’ indicates a point of 2% increase. ‘PIP max’ was set at the point that marks a drop in signal at least 70% as steep as the maximum slope observed in each individual cell. ‘PIP mid’ was set at the point corresponding to 50% difference between ‘PIP min’ and ‘max’. Geminin_1-110_ ‘rise’ was defined as the first point that shows increase of either 3% over minimum signal or 5 arbitrary intensity units.

### PIP signal after NCS treatment

An asynchronous population of PIP-FUCCI U2OS cells was imaged for 20 h before NCS treatment. Cells in G1 or G2 phases at the moment of NCS addition, as classified by PCNA pattern, were selected for analysis (tracking, segmentation of nuclear ROIs followed by PIP intensity measurement). The drop in signal was calculated based on the intensity traces of PIP 2 h after NCS treatment. In G1 cells, the drop was calculated as a change between mean signal before the stimulation and the signal measured 2 h after NCS treatment. In G2 cells it was assumed that PIP signal increases linearly within G2 phase. Signals from cells that showed at least three G2 measurements before NCS treatment were fitted to a linear function, and the parameters obtained were used to predict the signal intensity 2 h after NCS addition. The loss of PIP signal was measured as the difference between this prediction value and the actual measurement. As a control for this approach we analyzed a population of untreated cells in the same way, except that a mock NCS point was chosen randomly for each individual cell.

## Acknowledgements

We thank Jeffrey Jones for managerial assistance, Dr. Sam Wolff for microscopy assistance, and Cere Poovey for technical assistance. We also thank Dr. Steven Cappell, Dr. Robert Duronio, Dr. Michael Emanuele, and the members of our labs for helpful discussions. This work was supported by grants from the NIH to J.G.C. (GM083024 and GM102413), G.D.G. (T32CA009156), J.C.L. (T32GM007040), and to J.E.P. (DP2-HD091800). Additional funding was provided by the W.M. Keck foundation to J.E.P. and J.G.C.

## Supplementary figures

**Figure S1.**
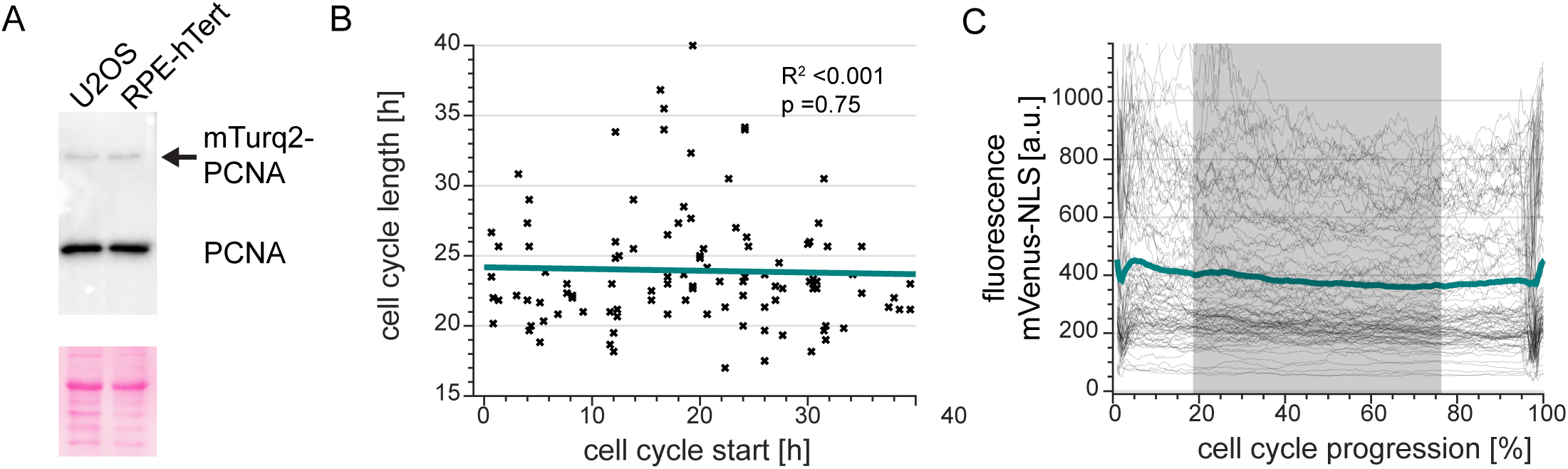
Total Cell cycle time does not vary during long imaging runs; relative PCNA expression. A. Whole cell lysates of U2OS and RPE-hTert cells expressing PIP-FUCCI reporters and mTurq2-PCNA were subjected to immunoblotting for endogenous (lower band) and ectopic (upper band marked with an arrow) PCNA protein levels. Lower panel is the corresponding portion of the blot stained for total protein with Ponceau S.
B. Analysis of cell cycle duration in the population of analyzed U2OS PIP-FUCCI cells born in different time points (x-axis) during a 64 h experiment. Teal line – best linear fit, n=107.
C. Warping (see Materials and Methods and main Fig.1D) of a constitutively stable reporter (mVenus-NLS expressed in U2OS cells) to detect any artefactual oscillations at the transition in and out of S phase. Small changes visible at the beginning and end of the cell cycle correspond to changes in the nuclear region area, release into the cytoplasm after nuclear envelope breakdown and subsequent packing of the construct into the nucleus after mitosis. Thin gray lines – individual cells (n=97), teal line – mean signal, gray rectangle – S phase as detected by PCNA pattern analysis.

**Figure S2.**
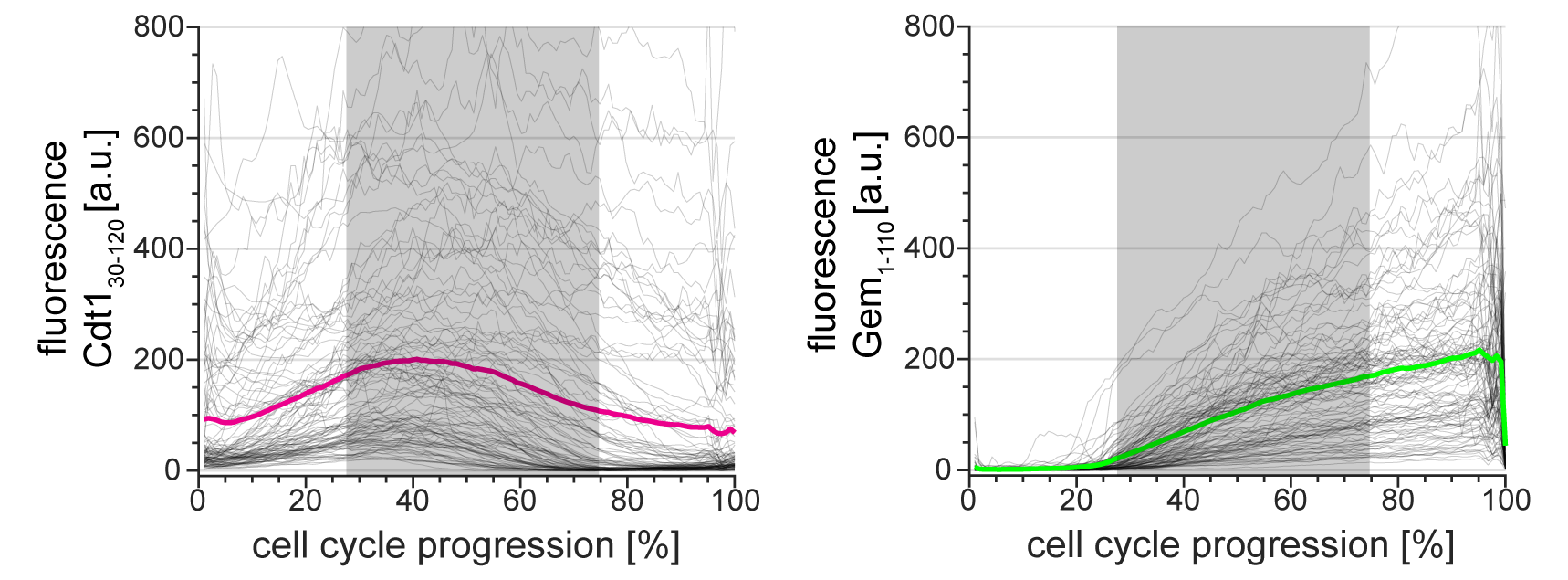
Cdt1_30-120_ dynamics in high-expressing FUCCI cells are dampened. Single cell traces (gray lines) and mean (red and green) of FUCCI reporter dynamics in a particularly bright clone of U2OS cell line (n=111). All traces were warped as in figure 1D; gray rectangles show the median S phase duration of this clone as detected based on PCNA variance.

**Figure S3.**
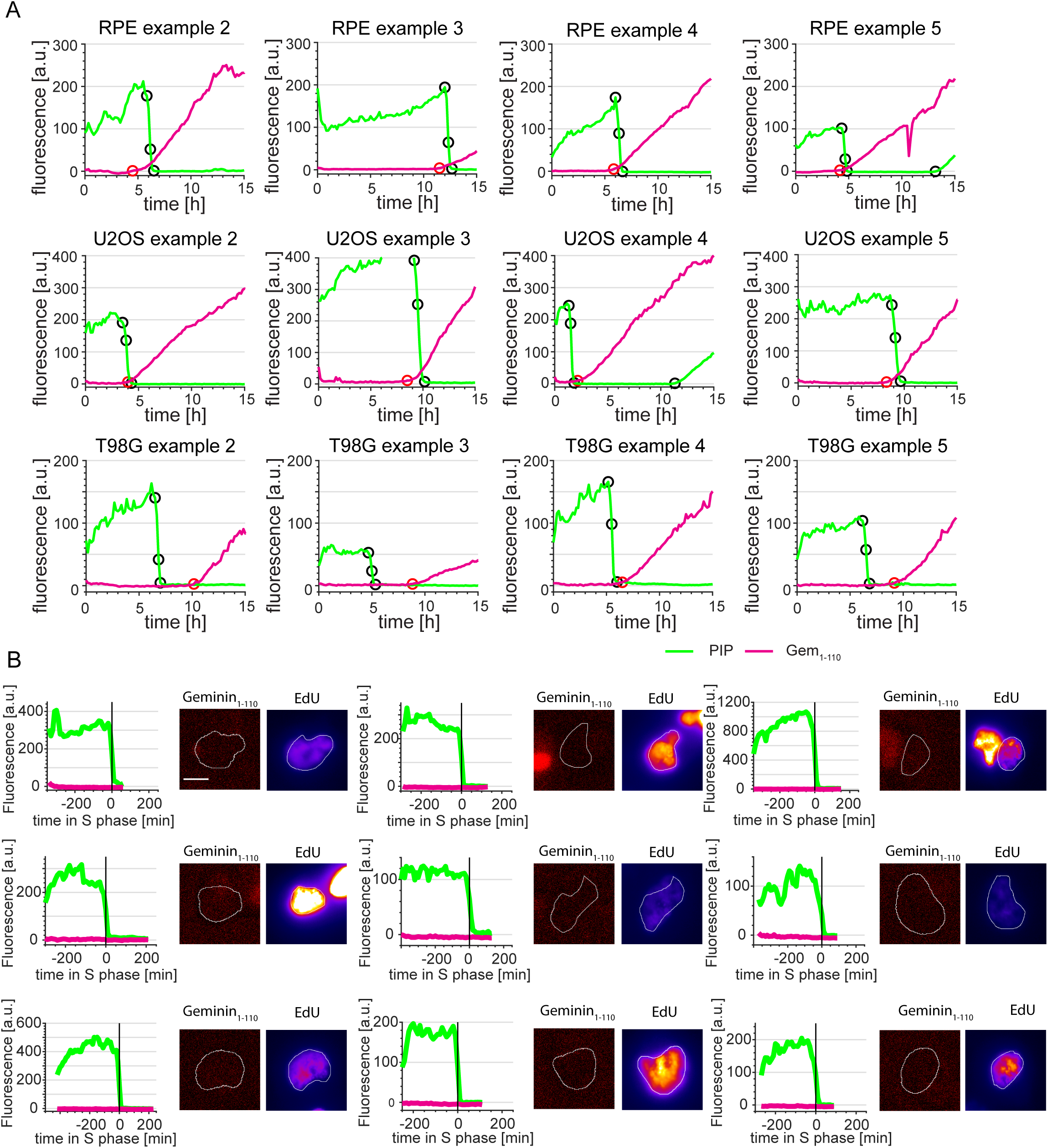
Gem_1-110_ dynamics in the three cell lines. A. Complementary to Figure 6A. Additional examples of RPE-hTert (upper row), U2OS (middle row) and T98G (lower row) cells transitioning from G1 to S phase with different temporal relationships between PIP-mVenus and mCherry-Gem_1-110_ signals.
B. Complementary to Figure 6C. Additional examples of early S phase T98G cells showing degradation of PIP-mVenus and EdU incorporation but no mCherry-Gem_1-110_ signal increase. EdU signal presented using ‘Fire’ look-up table (Fiji, black – min, yellow - max).

**Figure S4.**
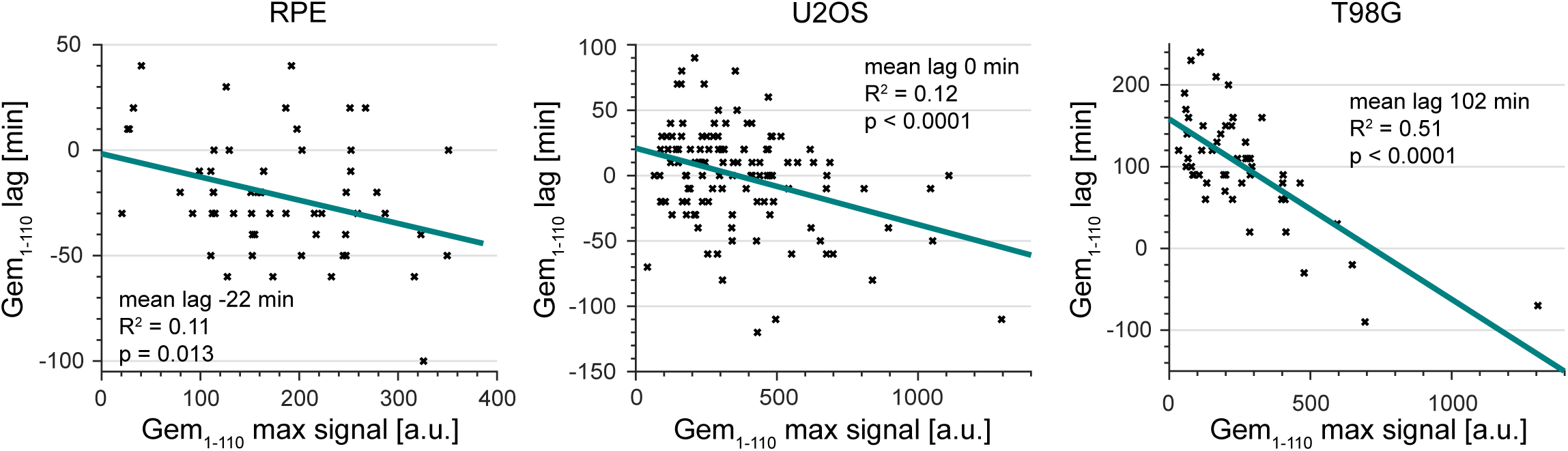
Gem_1-110_ increase is detected earlier in brighter cells. Correlation between maximum signal of mCherry-Gem_1-110_ and Gem_1-110_ lag in RPE-hTert, U2OS and T98G cells (RPE-hTert n=56, U2OS n=120, T98G n=51).

**Figure S5.**
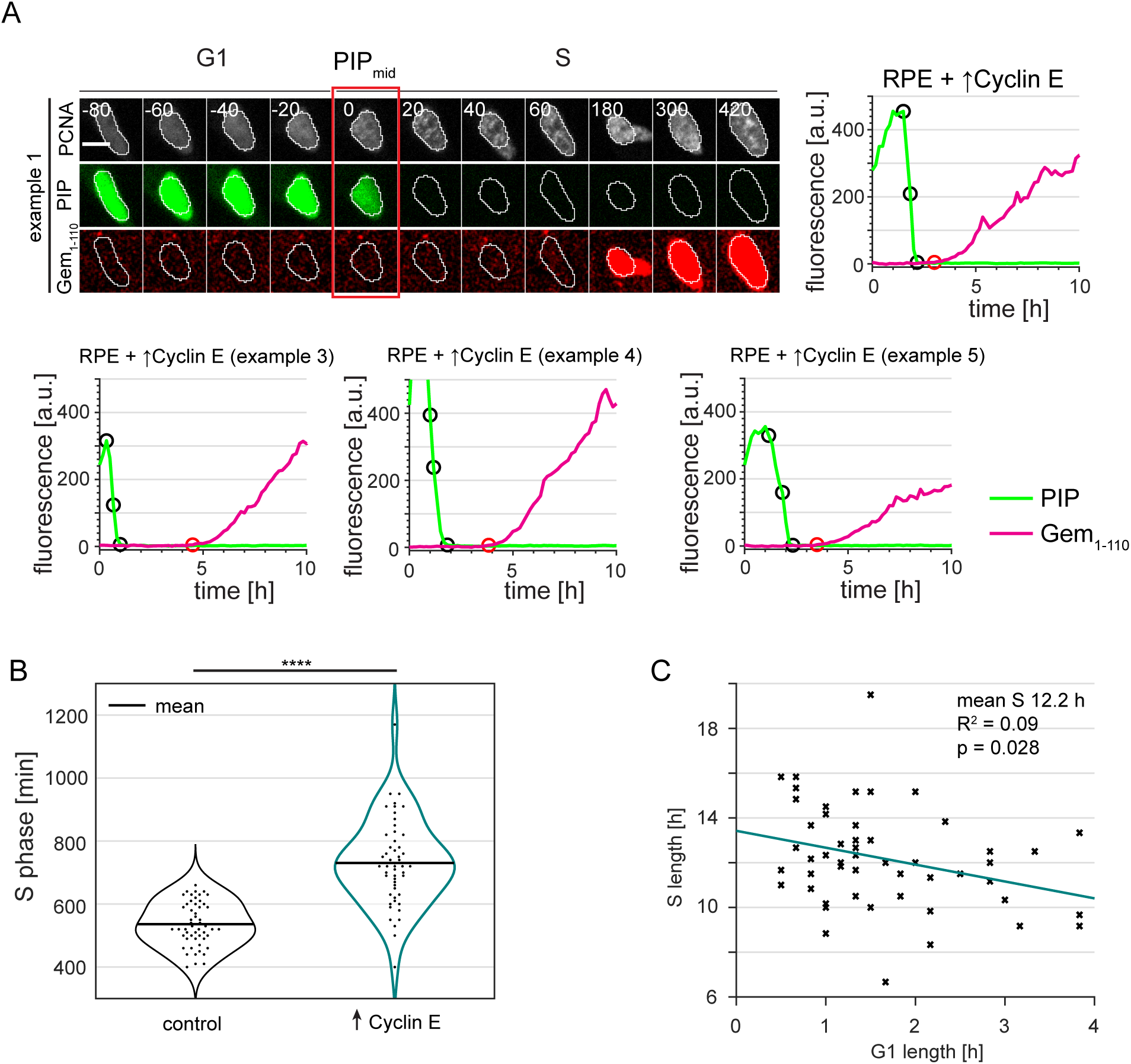
Gem_1-110_ dynamics in RPE-hTert cells overproducing cyclin E. A. Complimentary to Figure 7A. Additional examples of PRE cells overexpressing Cyclin E cells transitioning from G1 to S phase showing delay in mCherry-Gem_1-110_ accumulation.
B. Distributions of S length in control RPE-hTert (n=52) and in their isogenic counterparts overproducing Cyclin E (n=53). Kolmogorov-Smirnov test, ^⋆⋆⋆⋆^ p<0.0001.
C. Correlation between G1 length and S length in RPE-hTert cells overexpressing Cyclin E (n=53).

**Figure S6.**
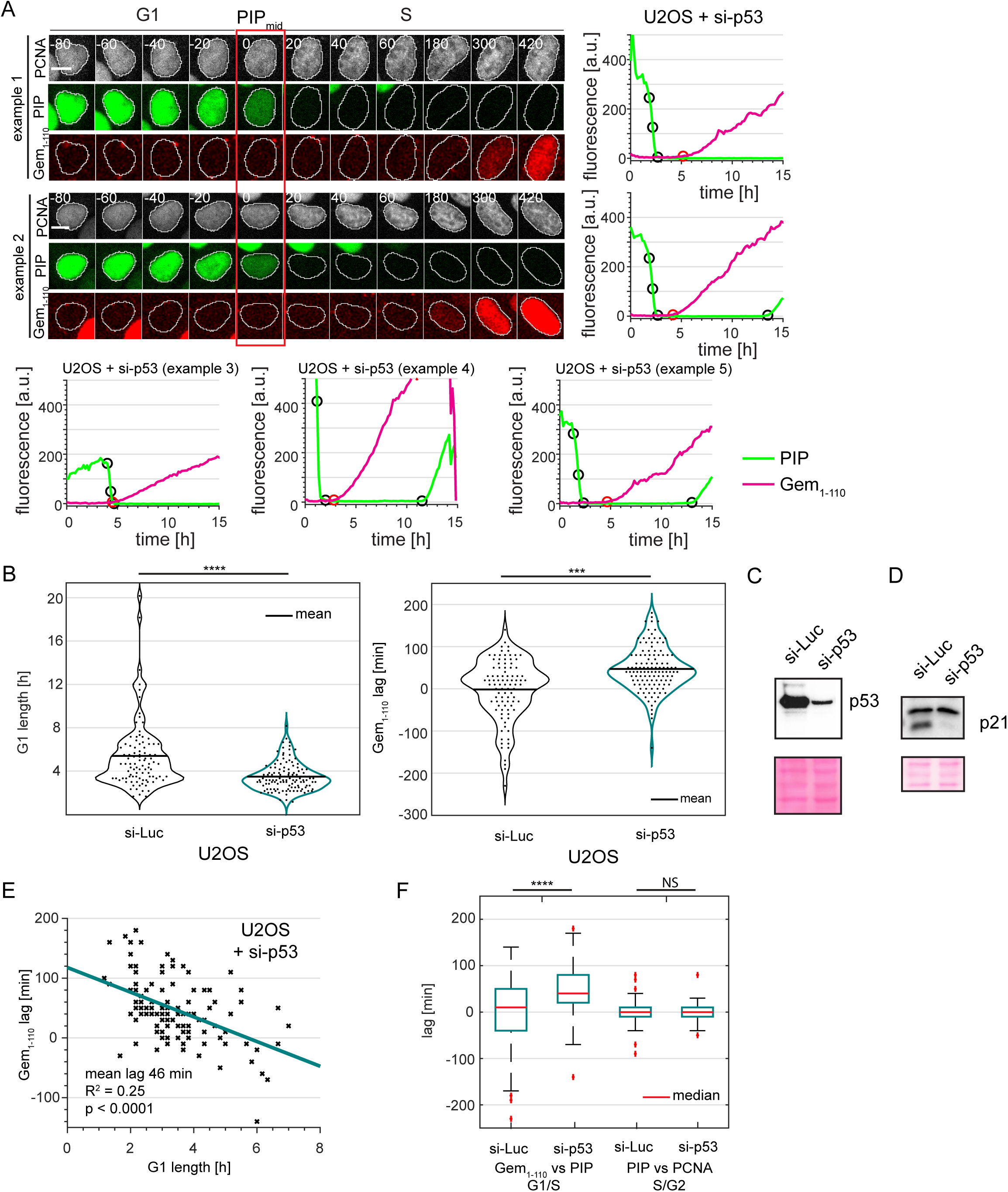
Gem_1-110_ dynamics in p53-depleted U2OS cells. A. Examples showing increased Gem_1-110_ lag in PIP-FUCCI U2OS cells transfected with siRNA targeting p53 prior to imaging. Numbers indicate minutes from the G1/S transition as measured by PIP-mVenus dynamics; scale bar 10 μm.
B. (Left) Distributions of G1 lengths in U2OS treated with si-Luciferase (n=97) and si-p53 (n=119). (Right) Distributions of mCherry-gem_1-110_ lag in the same cells as the left panel. Positive lag indicates mCherry-gem_1-110_ accumulation started after S phase. Kolmogorov-Smirnov test, ^⋆⋆⋆^ p<0.001, ^⋆⋆⋆⋆^ p<0.0001.
C. Parallel cultures of PIP-FUCCI and mTurq2-PCNA U2OS cells transfected with si-p53 or a control siRNA at the same time as the cells in Figure 6D. After 48 hours whole cell lysates were immunoblotted for endogenous p53. Lower panel is the corresponding portion of the blot stained for total protein with Ponceau S.
D. The same lysates in C. immunoblotted for endogenous p21.
E. Correlation between G1 length and Gem_1-110_ lag in p53-depleted U2OS cells (n=119).
F. Distributions of lag in detection of mCherry-Gem_1-110_ at the beginning of S phase (as indicated by PIP_mid_) and PIP-mVenus at the beginning of G2 phase (as indicated by PCNA pattern) in U2OS cells treated with si-Luciferase (n=97) and si p53 (n=119). Kolmogorov-Smirnov test, ^⋆⋆⋆⋆^ p<0.0001.

## Supplementary movies

### Movie 1

A single U2OS cells expressing PIP-FUCCI and PCNA throughout the cell cycle with a graph showing quantification of the signals (green: PIP-mVenus, magenta: mCherry-Gem1-110, black – variance of PCNA signal). Scale bar, 10 μm.

### Movie 2

RPE-hTert cells expressing PIP-FUCCI (green: PIP-mVenus, red: mCherry-Gem1-110) and PCNA (not shown) in asynchronous population. Gray color shows DIC signal, scale bar, 50 μm.

### Movie 3

RPE-hTert (control, no Dox) cells expressing PIP-FUCCI (green: PIP-mVenus, red: mCherry-Gem1-110 and gray: PCNA) in asynchronous population. Scale bar, 50 μm. Please note G1 length of around 8 h and cells immediately increasing mCherry-Gem1-110 signal upon entering S phase.

### Movie 4

RPE-hTert overexpressing Cyclin E (5 ng/ml Dox) with PIP-FUCCI (green: PIP-mVenus, red: mCherry-Gem1-110 and gray: PCNA) in asynchronous population. Scale bar, 50 μm. Please note G1 length shorter than 2 h and cells entering S phase without mCherry-Gem1-110 present.

## References

1. Davis DM, Purvis JE. Computational analysis of signaling patterns in single cells. Semin Cell Dev Biol. 2015;37:35–43.

2. Matson JP, Cook JG. Cell cycle proliferation decisions: the impact of single cell analyses. FEBS J. 2017;284(3):362–75.

3. Spencer SL, Cappell SD, Tsai FC, Overton KW, Wang CL, Meyer T. The Proliferation-Quiescence Decision Is Controlled by a Bifurcation in CDK2 Activity at Mitotic Exit. Cell. 2013.

4. Sakaue-Sawano A, Kurokawa H, Morimura T, Hanyu A, Hama H, Osawa H, et al. Visualizing spatiotemporal dynamics of multicellular cell-cycle progression. Cell. 2008; 132(3):487–98.

5. Nishitani H, Sugimoto N, Roukos V, Nakanishi Y, Saijo M, Obuse C, et al. Two E3 ubiquitin ligases, SCF-Skp2 and DDB1-Cul4, target human Cdt1 for proteolysis. EMBO J. 2006;25(5):1126–36.

6. McGarry TJ, Kirschner MW. Geminin, an inhibitor of DNA replication, is degraded during mitosis. Cell. 1998;93(6):1043–53.

7. Fukuhara S, Zhang J, Yuge S, Ando K, Wakayama Y, Sakaue-Sawano A, et al. Visualizing the cell-cycle progression of endothelial cells in zebrafish. Dev Biol. 2014;393(1):10–23.

8. Sladitschek HL, Neveu PA. MXS-Chaining: A Highly Efficient Cloning Platform for Imaging and Flow Cytometry Approaches in Mammalian Systems. PLoS One. 2015; 10(4):e0124958.

9. Sugiyama M, Sakaue-Sawano A, Iimura T, Fukami K, Kitaguchi T, Kawakami K, et al. Illuminating cell-cycle progression in the developing zebrafish embryo. Proc Natl Acad Sci U S A. 2009; 106(49):20812–7.

10. Zielke N, Korzelius J, van Straaten M, Bender K, Schuhknecht GF, Dutta D, et al. Fly-FUCCI: A versatile tool for studying cell proliferation in complex tissues. Cell reports. 2014;7(2):588–98.

11. Zielke N, Edgar BA. FUCCI sensors: powerful new tools for analysis of cell proliferation. Wiley Interdiscip Rev Dev Biol. 2015;4(5):469–87.

12. Sakaue-Sawano A, Yo M, Komatsu N, Hiratsuka T, Kogure T, Hoshida T, et al. Genetically Encoded Tools for Optical Dissection of the Mammalian Cell Cycle. Mol Cell. 2017.

13. Leonhardt H, Rahn HP, Weinzierl P, Sporbert A, Cremer T, Zink D, et al. Dynamics of DNA replication factories in living cells. J Cell Biol. 2000;149(2):271–80.

14. Burgess A, Lorca T, Castro A. Quantitative live imaging of endogenous DNA replication in mammalian cells. PLoS One. 2012;7(9):e45726.

15. Zerjatke T, Gak IA, Kirova D, Fuhrmann M, Daniel K, Gonciarz M, et al. Quantitative Cell Cycle Analysis Based on an Endogenous All-in-One Reporter for Cell Tracking and Classification. Cell reports. 2017;19(9):1953–66.

16. Madsen P, Celis JE. S-phase patterns of cyclin (PCNA) antigen staining resemble topographical patterns of DNA synthesis. A role for cyclin in DNA replication? FEBS Lett. 1985;193(1):5–11.

17. Kennedy BK, Barbie DA, Classon M, Dyson N, Harlow E. Nuclear organization of DNA replication in primary mammalian cells. Genes Dev. 2000;14(22):2855–68.

18. Chao HX, Poovey CE, Privette AA, Grant GD, Chao HY, Cook JG, et al. Orchestration of DNA Damage Checkpoint Dynamics across the Human Cell Cycle. Cell Syst. 2017;5(5):445–59 e5.

19. Wilson KA, Elefanty AG, Stanley EG, Gilbert DM. Spatio-temporal re-organization of replication foci accompanies replication domain consolidation during human pluripotent stem cell lineage specification. Cell Cycle. 2016:0.

20. Overton KW, Spencer SL, Noderer WL, Meyer T, Wang CL. Basal p21 controls population heterogeneity in cycling and quiescent cell cycle states. Proc Natl Acad Sci U S A. 2014;111(41):E4386–93.

21. Spiller DG, Wood CD, Rand DA, White MR. Measurement of single-cell dynamics. Nature. 2010;465(7299):736–45.

22. Chiorino G, Metz JA, Tomasoni D, Ubezio P. Desynchronization rate in cell populations: mathematical modeling and experimental data. J Theor Biol. 2001;208(2): 185–99.

23. Snijder B, Pelkmans L. Origins of regulated cell-to-cell variability. Nat Rev Mol Cell Biol. 2011; 12(2): 119–25.

24. Dueck H, Eberwine J, Kim J. Variation is function: Are single cell differences functionally important?: Testing the hypothesis that single cell variation is required for aggregate function. Bioessays. 2016;38(2):172–80.

25. Darzynkiewicz Z, Crissman H, Traganos F, Steinkamp J. Cell heterogeneity during the cell cycle. J Cell Physiol. 1982;113(3):465–74.

26. Takeda DY, Parvin JD, Dutta A. Degradation of Cdt1 during S phase is Skp2-independent and is required for efficient progression of mammalian cells through S phase. J Biol Chem. 2005;280(24):23416–23.

27. Hornbeck PV, Zhang B, Murray B, Kornhauser JM, Latham V, Skrzypek E. PhosphoSitePlus, 2014: mutations, PTMs and recalibrations. Nucleic Acids Res. 2015;43(Database issue):D512–20.

28. Coleman KE, Grant GD, Haggerty RA, Brantley K, Shibata E, Workman BD, et al. Sequential replication-coupled destruction at G1/S ensures genome stability. Genes Dev. 2015;29(16):1734–46.

29. Kim JH, Lee SR, Li LH, Park HJ, Park JH, Lee KY, et al. High cleavage efficiency of a 2A peptide derived from porcine teschovirus-1 in human cell lines, zebrafish and mice. PLoS One. 2011;6(4):e18556.

30. Nishitani H, Shiomi Y, Iida H, Michishita M, Takami T, Tsurimoto T. CDK inhibitor p21 is degraded by a proliferating cell nuclear antigen-coupled Cul4-DDB1Cdt2 pathway during S phase and after UV irradiation. J Biol Chem. 2008;283(43):29045–52.

31. Abbas T, Sivaprasad U, Terai K, Amador V, Pagano M, Dutta A. PCNA-dependent regulation of p21 ubiquitylation and degradation via the CRL4Cdt2 ubiquitin ligase complex. Genes Dev. 2008;22(18):2496–506.

32. Stuart SA, Wang JY. Ionizing radiation induces ATM-independent degradation of p21Cip1 in transformed cells. J Biol Chem. 2009;284(22):15061–70.

33. Zhang S, Zhao H, Darzynkiewicz Z, Zhou P, Zhang Z, Lee EY, et al. A novel function of CRL4(Cdt2): regulation of the subunit structure of DNA polymerase delta in response to DNA damage and during the S phase. J Biol Chem. 2013;288(41):29550–61.

34. Aleksandrov R, Dotchev A, Poser I, Krastev D, Georgiev G, Panova G, et al. Protein Dynamics in Complex DNA Lesions. Mol Cell. 2018;69(6):1046–61 e5.

35. Stivala LA, Prosperi E, Rossi R, Bianchi L. Involvement of proliferating cell nuclear antigen in DNA repair after damage induced by genotoxic agents in human fibroblasts. Carcinogenesis. 1993;14(12):2569–73.

36. Li R, Waga S, Hannon GJ, Beach D, Stillman B. Differential effects by the p21 CDK inhibitor on PCNA-dependent DNA replication and repair. Nature. 1994;371(6497):534–7.

37. Shivji KK, Kenny MK, Wood RD. Proliferating cell nuclear antigen is required for DNA excision repair. Cell. 1992;69(2):367–74.

38. Arias EE, Walter JC. PCNA functions as a molecular platform to trigger Cdt1 destruction and prevent re-replication. Nat Cell Biol. 2006;8(1):84–90.

39. Hu J, Xiong Y. An evolutionarily conserved function of proliferating cell nuclear antigen for Cdt1 degradation by the Cul4-Ddb1 ubiquitin ligase in response to DNA damage. J Biol Chem. 2006;281 (7):3753–6.

40. Rizzardi LF, Coleman KE, Varma D, Matson JP, Oh S, Cook JG. CDK1-dependent inhibition of the E3 ubiquitin ligase CRL4CDT2 ensures robust transition from S Phase to Mitosis. J Biol Chem. 2015;290(1):556–67.

41. Nukina K, Hayashi A, Shiomi Y, Sugasawa K, Ohtsubo M, Nishitani H. Mutations at multiple CDK phosphorylation consensus sites on Cdt2 increase the affinity of CRL4(Cdt2) for PCNA and its ubiquitination activity in S phase. Genes Cells. 2018.

42. Johmura Y, Shimada M, Misaki T, Naiki-Ito A, Miyoshi H, Motoyama N, et al. Necessary and sufficient role for a mitosis skip in senescence induction. Mol Cell. 2014;55(1):73–84.

43. Krenning L, Feringa FM, Shaltiel IA, van den Berg J, Medema RH. Transient activation of p53 in G2 phase is sufficient to induce senescence. Mol Cell. 2014;55(1):59–72.

44. Suzuki M, Yamauchi M, Oka Y, Suzuki K, Yamashita S. Live-cell imaging visualizes frequent mitotic skipping during senescence-like growth arrest in mammary carcinoma cells exposed to ionizing radiation. Int J Radiat Oncol Biol Phys. 2012;83(2):e241–50.

45. Mendez J, Stillman B. Chromatin association of human origin recognition complex, cdc6, and minichromosome maintenance proteins during the cell cycle: assembly of prereplication complexes in late mitosis. Mol Cell Biol. 2000;20(22):8602–12.

46. Vaziri C, Saxena S, Jeon Y, Lee C, Murata K, Machida Y, et al. A p53-dependent checkpoint pathway prevents rereplication. Mol Cell. 2003;11(4):997–1008.

47. Melixetian M, Ballabeni A, Masiero L, Gasparini P, Zamponi R, Bartek J, et al. Loss of Geminin induces rereplication in the presence of functional p53. J Cell Biol. 2004;165(4):473–82.

48. Davidson IF, Li A, Blow JJ. Deregulated replication licensing causes DNA fragmentation consistent with head-to-tail fork collision. Mol Cell. 2006;24(3):433–43.

49. Hall JR, Lee HO, Bunker BD, Dorn ES, Rogers GC, Duronio RJ, et al. Cdt1 and Cdc6 are destabilized by rereplication-induced DNA damage. J Biol Chem. 2008;283(37):25356–63.

50. Klotz-Noack K, McIntosh D, Schurch N, Pratt N, Blow JJ. Re-replication induced by geminin depletion occurs from G2 and is enhanced by checkpoint activation. J Cell Sci. 2012;125(Pt 10):2436–45.

51. Truong LN, Wu X. Prevention of DNA re-replication in eukaryotic cells. J Mol Cell Biol. 2011;3(1):13–22.

52. Stein GH. T98G: an anchorage-independent human tumor cell line that exhibits stationary phase G1 arrest in vitro. J Cell Physiol. 1979;99(1):43–54.

53. Cappell SD, Chung M, Jaimovich A, Spencer SL, Meyer T. Irreversible APC(Cdh1) Inactivation Underlies the Point of No Return for Cell-Cycle Entry. Cell. 2016;166(1):167–80.

54. Merzlyak EM, Goedhart J, Shcherbo D, Bulina ME, Shcheglov AS, Fradkov AF, et al. Bright monomeric red fluorescent protein with an extended fluorescence lifetime. Nat Methods. 2007;4(7):555–7.

55. Ekholm-Reed S, Mendez J, Tedesco D, Zetterberg A, Stillman B, Reed SI. Deregulation of cyclin E in human cells interferes with prereplication complex assembly. J Cell Biol. 2004;165(6):789–800.

56. Resnitzky D, Gossen M, Bujard H, Reed SI. Acceleration of the G1/S phase transition by expression of cyclins D1 and E with an inducible system. Mol Cell Biol. 1994;14(3):1669–79.

57. Bashir T, Dorrello NV, Amador V, Guardavaccaro D, Pagano M. Control of the SCF(Skp2-Cks1) ubiquitin ligase by the APC/C(Cdh1) ubiquitin ligase. Nature. 2004;428(6979):190–3.

58. Wei W, Ayad NG, Wan Y, Zhang GJ, Kirschner MW, Kaelin WG, Jr. Degradation of the SCF component Skp2 in cell-cycle phase G1 by the anaphase-promoting complex. Nature. 2004;428(6979):194–8.

59. Yin K, Ueda M, Takagi H, Kajihara T, Sugamata Aki S, Nobusawa T, et al. A dual-color marker system for in vivo visualization of cell cycle progression in Arabidopsis. Plant J. 2014;80(3):541–52.

60. Bajar BT, Lam AJ, Badiee RK, Oh YH, Chu J, Zhou XX, et al. Fluorescent indicators for simultaneous reporting of all four cell cycle phases. Nat Methods. 2016;13(12):993–6.

61. Ge XQ, Blow JJ. Chk1 inhibits replication factory activation but allows dormant origin firing in existing factories. J Cell Biol. 2010;191(7):1285–97.

62. Chandrasekaran S, Tan TX, Hall JR, Cook JG. Stress-stimulated mitogen-activated protein kinases control the stability and activity of the Cdt1 DNA replication licensing factor. Mol Cell Biol. 2011; 31(22):4405–16.

63. Clijsters L, Ogink J, Wolthuis R. The spindle checkpoint, APC/C(Cdc20), and APC/C(Cdh1) play distinct roles in connecting mitosis to S phase. J Cell Biol. 2013;201(7):1013–26.

64. Ballabeni A, Melixetian M, Zamponi R, Masiero L, Marinoni F, Helin K. Human geminin promotes pre-RC formation and DNA replication by stabilizing CDT1 in mitosis. EMBO J. 2004;23(15):3122–32.

65. Ballabeni A, Zamponi R, Moore JK, Helin K, Kirschner MW. Geminin deploys multiple mechanisms to regulate Cdt1 before cell division thus ensuring the proper execution of DNA replication. Proc Natl Acad Sci U S A. 2013;110(30):E2848–53.

66. Coulombe P, Gregoire D, Tsanov N, Mechali M. A spontaneous Cdt1 mutation in 129 mouse strains reveals a regulatory domain restraining replication licensing. Nat Commun. 2013;4:2065.

67. Hahn AT, Jones JT, Meyer T. Quantitative analysis of cell cycle phase durations and PC12 differentiation using fluorescent biosensors. Cell Cycle. 2009;8(7):1044–52.

68. Yao G, Lee TJ, Mori S, Nevins JR, You L. A bistable Rb-E2F switch underlies the restriction point. Nat Cell Biol. 2008;10(4):476–82.

69. Lukas C, Sorensen CS, Kramer E, Santoni-Rugiu E, Lindeneg C, Peters JM, et al. Accumulation of cyclin B1 requires E2F and cyclin-A-dependent rearrangement of the anaphase-promoting complex. Nature. 1999;401(6755):815–8.

70. Choudhury R, Bonacci T, Arceci A, Lahiri D, Mills CA, Kernan JL, et al. APC/C and SCF(cyclin F) Constitute a Reciprocal Feedback Circuit Controlling S-Phase Entry. Cell reports. 2016;16(12):3359–72.

71. Fukushima H, Ogura K, Wan L, Lu Y, Li V, Gao D, et al. SCF-mediated Cdh1 degradation defines a negative feedback system that coordinates cell-cycle progression. Cell reports. 2013;4(4):803–16.

72. Yuan X, Srividhya J, De Luca T, Lee JH, Pomerening JR. Uncovering the role of APC-Cdh1 in generating the dynamics of S-phase onset. Mol Biol Cell. 2014;25(4):441–56.

73. Matson JP, Dumitru R, Coryell P, Baxley RM, Chen W, Twaroski K, et al. Rapid DNA replication origin licensing protects stem cell pluripotency. Elife. 2017;6.

74. Schindelin J, Arganda-Carreras I, Frise E, Kaynig V, Longair M, Pietzsch T, et al. Fiji: an open-source platform for biological-image analysis. Nat Methods. 2012;9(7):676–82.

